# Structural basis for TORC2 activation

**DOI:** 10.1101/2025.07.02.662585

**Authors:** Luoming Zou, Maria G. Tettamanti, Ariane Bergmann, Robbie Loewith, Lucas Tafur

## Abstract

The Target of Rapamycin Complex 2 (TORC2) is a central node in signaling feedback loops serving to maintain biophysical homeostasis of the plasma membrane (PM). How TORC2 is regulated by mechanical perturbation of the PM is not well understood. To address this, we determined the cryo-electron microscopy structure of endogenous yeast TORC2 to up to 2.2 Å resolution. Our model refines the position and interactions of TORC2-specific subunits, providing a structural basis for the differential assembly of Tor2 into TORC2. Furthermore, we observe the insertion of the pleckstrin-homology domain of the Avo1 subunit into the Tor2 active site, providing a regulatory mechanism by phosphoinositides. Structure-guided functional experiments reveal a potential TORC2 membrane binding surface and a positively charged pocket in the Avo3 subunit that is necessary for TORC2 activation. Collectively, our data suggest that signaling phosphoinositides activate TORC2 by membrane-induced structural rearrangements via concerted action of conserved regulatory subunits.

## Introduction

The Target Of Rapamycin (TOR) is a highly conserved serine/threonine kinase that controls cell growth and metabolism across eukaryotes^1^. In yeast and mammals, TOR forms part of two multi-component complexes, TORC1 and TORC2, each with distinct localization and downstream targets^2^. In yeast, TORC2 is localized to the plasma membrane (PM), regulating signaling pathways that act in complex feedback loops to maintain the biophysical homeostasis of the PM^3–6^. In mammals, mTORC2 also appears to be localized to the PM, although not exclusively^7^. In contrast to (m)TORC1, (m)TORC2 is rapamycin insensitive^2,8^, and hence, its regulation has been less straightforward to characterize.

TORC2 is composed of two copies of Tor2, Lst8, Avo1, Avo2, Avo3, and Bit61 or its paralog Bit2^9^ (Figure 1A). Avo1 appears to play a particularly important role in TORC2 function, as it mediates substrate recruitment via its CRIM (conserved region in the middle) domain^10^. Avo1 also contains a Ras-binding domain (RBD)^11,12^ and a PH (pleckstrin-homology) domain in the C-terminus, which binds to regulatory phosphoinositides (PtdIns)^13^. The PH domain of the mammalian Avo1 ortholog, mSin1, has been proposed to interact with the mTOR kinase domain to suppress mTORC2 signaling^14^. Avo3 is the TORC2 counterpart of Kog1 in TORC1, binding in a mutually exclusive manner to the TOR kinase^2,9^. Importantly, the Avo3 C-terminal domain interacts with the TOR kinase domain and blocks the binding of FKBP12-rapamycin, explaining why TORC2 is insensitive to rapamycin^9,15^. Avo3 also appears to play an essential role in membrane recruitment of TORC2^16^. The function of Bit61 and Bit2 is much less understood; no phenotype has been reported for *bit61 bit2* mutants. It has been proposed that these subunits are orthologs of the equally mysterious Protor1/2 subunits of mTORC2 due to the shared presence of an HbrB domain^17^, although some debate remains^13^. Finally, Avo2, which is non-essential and apparently not conserved in mTORC2, has been proposed to stimulate the catalytic function of TORC2^18^.

**Figure 1.**
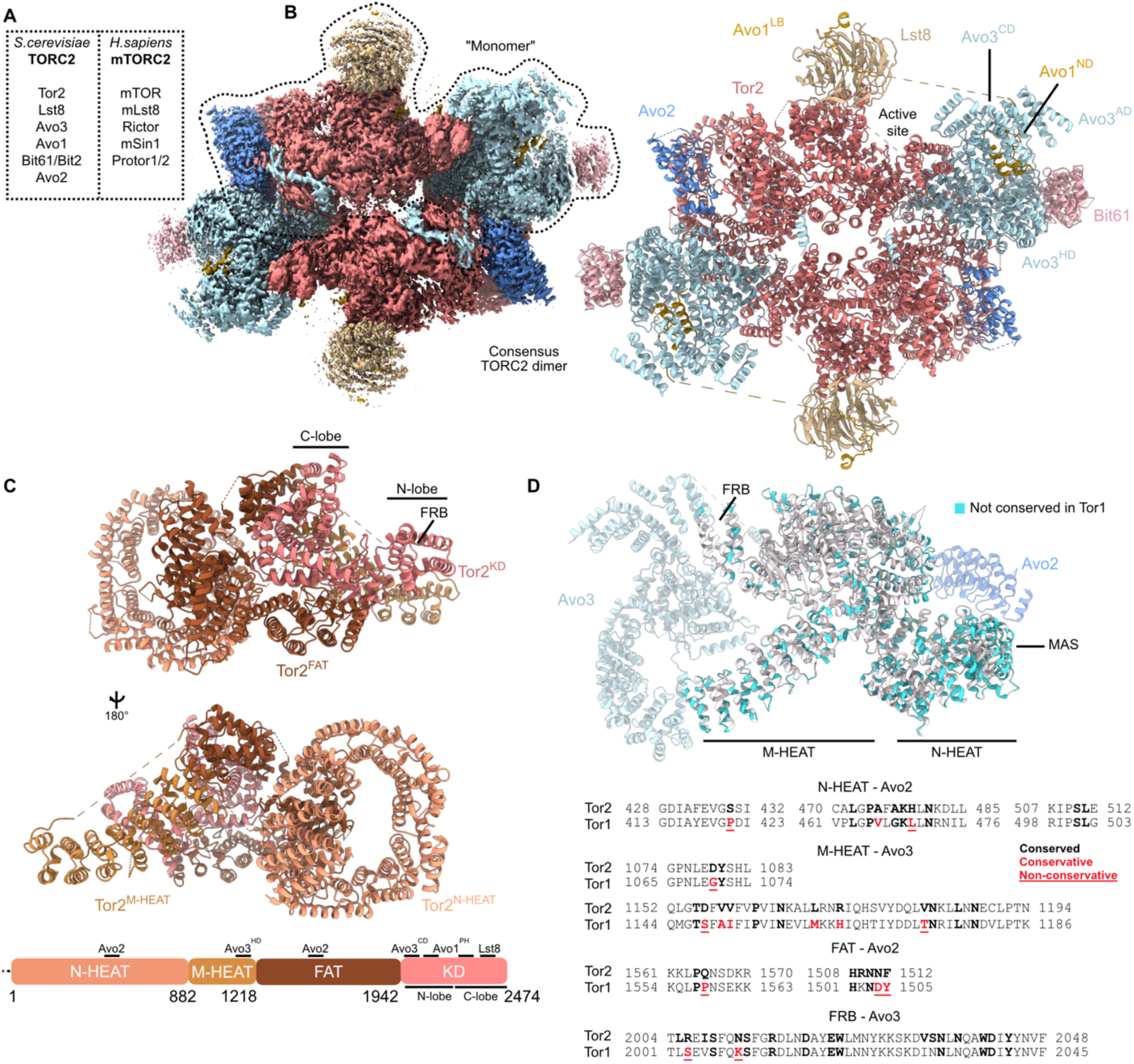
High resolution cryo-EM structure of endogenous TORC2. **A.** Subunits forming TORC2 in yeast and mTORC2 in humans. **B.** Cryo-EM density of “consensus” TORC2 dimer, indicating the monomer used for focused classifications (left). Overall TORC2 dimer model, indicating one copy of each subunit in the monomer (right). **C.** Structure and domain organization of the Tor2 kinase. **D.** Conservation of amino acids between Tor1 and Tor2. Residues that interact with Avo2 and Avo3 are shown in bold black, those replaced by chemically similar amino acids in Tor1 are shown in bold red, and non-conservative changes in Tor1 are shown in underlined bold red. See also Figures S1, S2, S3, S4 and S5.

There has been continuing improvement in the structural biology of (m)TORC2. Initial low resolution cryo-electron microscopy (cryo-EM) structures of yeast TORC2 established the molecular basis for the rapamycin insensitivity of the complex and showed that mTORC1 and TORC2 had a similar overall topology^9,15^. Higher resolution structures of recombinant mTORC2 later confirmed these findings and resolved the interaction between subunits in greater detail^19,20^. However, major questions about the regulation of (m)TORC2 activity remain. Some authors have proposed that activation of mTORC2 requires only substrate recruitment to the PM without any significant conformational change in the complex^7^, while others argue for a PtdIns-mediated conformational change that evicts the PH domain of mSin1 from the active site to enable substrate access^14^. Although intriguing, this model lacks structural evidence as the PH domain of Avo1/mSin1 is unresolved in available TORC2/mTORC2 structures and it is not known how (m)TORC2 interacts with the PM. Furthermore, how Bit61/Protor1 is integrated into the complex and if it affects the “core” TORC2 structure is not known. Thus, to better understand the regulation of (m)TORC2 activity, we set out to determine a high-resolution and more complete structure of yeast TORC2.

## Results

### High-resolution cryo-EM structure of endogenous TORC2

To obtain a sample suitable for high-resolution cryo-EM, we optimized our previous purification protocol and sample preparation extensively, obtaining stable high-quality TORC2 (STAR Methods). Data were collected on a 300 kV Titan Krios equipped with a Falcon 4 direct electron detector and a Selectris X energy filter. After sorting out particles using 2D classification and Heterogeneous Refinement jobs, we obtained a 2.4 Å resolution overall “consensus” reconstruction of the TORC2 dimer (Figures 1B, S1 and S2). Using a series of focused masks to resolve different parts of the complex, we achieved reconstructions ranging from 2.2 to 2.5 Å resolution, which allowed us to build a model of all TORC2 subunits (Tor2, Lst8, Avo1, Avo2, Avo3, Bit61), including well-resolved inositol hexaphosphate (IP6) bound to Tor2 (Figures 1B, S2, S3 and Table 1). To our knowledge, these represent the highest resolution cryo-EM reconstructions obtained for any (m)TOR complex (or component of the pathway) to date. As expected^9^, the overall structure of TORC2 is highly similar to mTORC2 (Figures 1B and S4A), forming a tighter core than TORC1 (in its inhibited oligomer, TOROID^21^) and mTORC1 due to closer packing of each Tor2 copy in the dimer (Figure S4B).

**Table 1.**
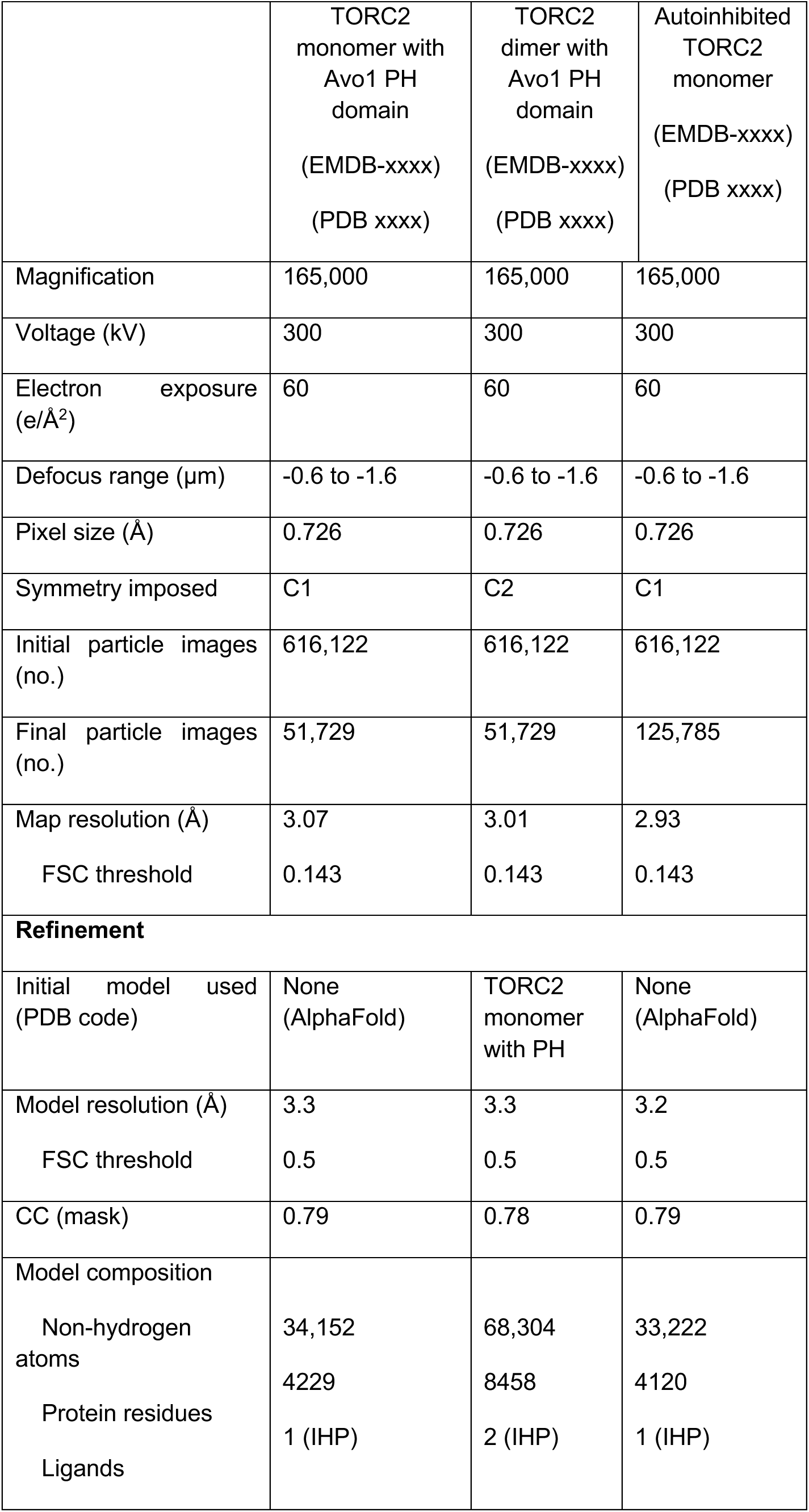
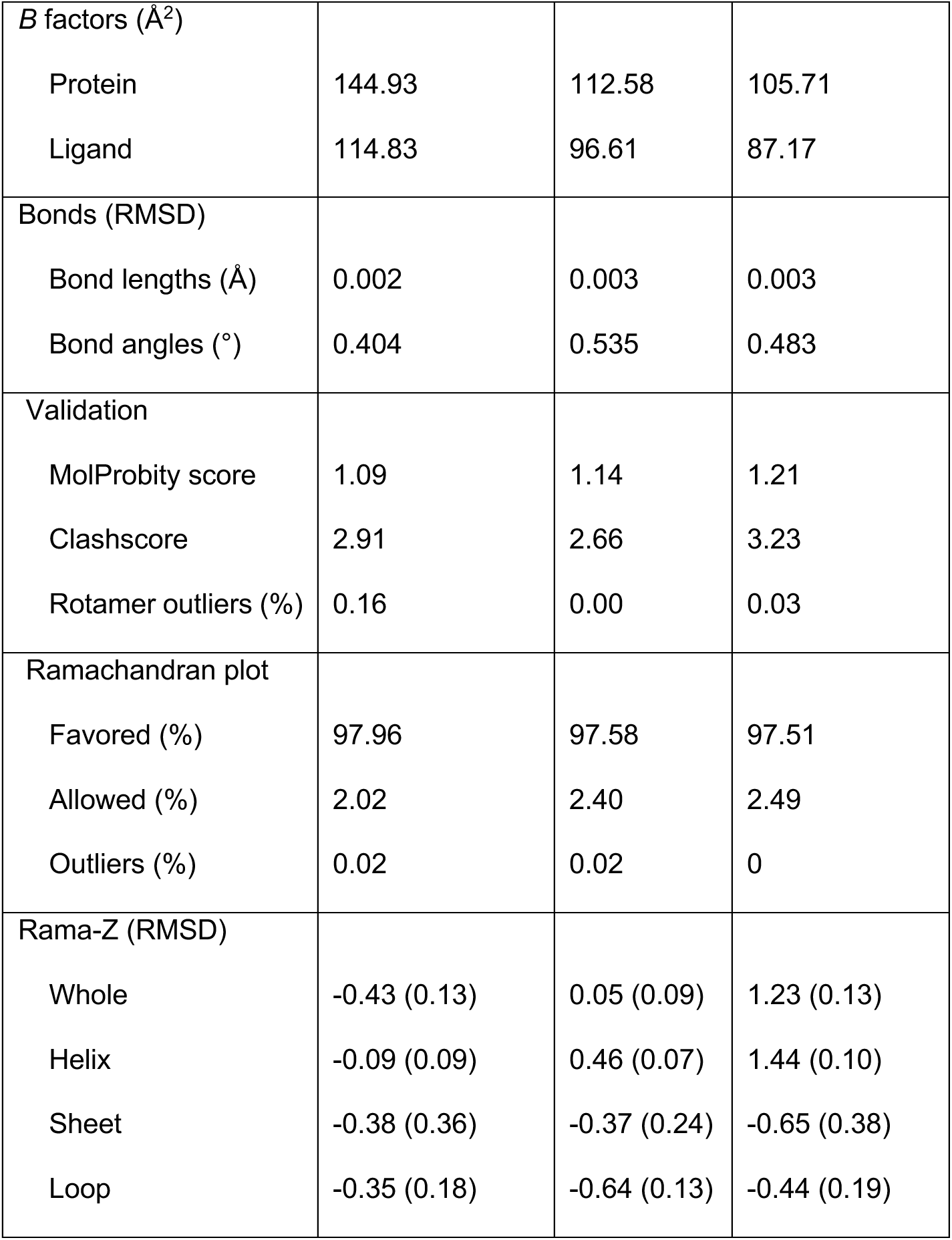
Cryo-EM data collection and refinement statistics.

During the analysis of the consensus map, we noticed poorly resolved density near the active site that was also visible in the previous low resolution cryo-EM map of TORC2^9^. To better resolve this region, we performed focused 3D classification on C2 symmetry-expanded particles (Figure S1). This revealed that most particles (77.6%) had extra density in this region. One small class displaying well-resolved density was further processed and refined to 3.07 Å resolution, revealing the position of the Avo1 PH domain (Avo1^PH^) (Figures S1 and S2). Likewise, we selected one class that clearly lacked any density in this region and obtained a 2.93 Å resolution reconstruction (Figures S1 and S2). We additionally noticed a weak density (that became more apparent at lower thresholds) in our reconstructions protruding from Lst8 that was also observed in the 2D classes from the previous 7.9 Å resolution cryo-EM map^9^. Using a similar strategy as for the Avo1^PH^ domain, we obtained a lower resolution volume that, based on its position along the sequence, we assign to the Avo1 CRIM domain (Avo1^CRIM^) (Figures S1 and S2). A similar low-resolution density next to mLst8 was previously observed in mTORC2 in the presence of its substrate Akt^19^. These different reconstructions are discussed in detail later.

### Specificity of Tor2 assembly in TORC2

The Tor2 kinase consists of four main regions: a flexible N-terminal HEAT repeat domain (N-HEAT, which is only resolved to low resolution), a middle HEAT repeat domain (M-HEAT), a FAT domain and a C-terminal Kinase Domain (KD) (Figure 1C). This architecture and structural organization are similar in Tor1^21^ and mTOR^19,22^.

In contrast to humans, which express a single mTOR kinase that forms part of both mTORC1 and mTORC2, yeast has two *TOR* genes, *TOR1* and *TOR2*^23^. Despite their similarity, only Tor2 is found in TORC2^2^, although the molecular basis for this has remained elusive. To better understand the reasons behind this difference, we first compared the conformation of Tor2 in TORC2 and TORC1 (as part of the TOROID^21^). Whereas the KD and FAT align relatively well, there is a relative shift in both HEAT domains (Figure S5A). This is likely due to TORC1 oligomerization, as the conformation of mTOR in mTORC1 and mTORC2 is similar (Figure S5B). As there is no structure of free TORC1, it is not clear if these changes are maintained when not forming TOROIDs.

A possible reason why Tor1 is unable to assemble TORC2 is because it cannot adopt the conformation of Tor2 in TORC2. This could be due to a lack of residues required for the interactions with Avo2 and Avo3, which are the subunits that contact Tor2 in the monomer, or because it has extra features that prevent these interactions. The AlphaFold prediction of Tor1 shows no obvious extensions in these regions that could prevent the interactions observed in TORC2 (Figure S5C). In contrast, when mapping the sequence conservation between Tor1 and Tor2 onto our structure, we observe several changes in residues that interact with Avo2 and Avo3 spread over all the domains of the kinase, suggesting that these differences drive the specificity of Tor2 (Figure 1D). The residues that contact Avo2 in the N-HEAT are located on the previously defined “major assembly specificity” (MAS) domain (residues 428-926), proposed to confer specificity for formation of either TORC1 or TORC2^24,25^. However, replacing the Tor2 MAS with the corresponding sequence of Tor1, while keeping the rest of Tor2 intact, allows the formation of TORC2^25^, suggesting that specificity is also driven by other regions. In agreement, our structure suggests that the specificity of Tor2 appears to be due to the contribution of multiple interactions along the protein rather than just a single interface, as suggested previously^13,24,25^.

### Avo3 organizes the TORC2 dimer

Focused refinement on Avo3 allowed us to obtain a 2.2 Å resolution map and thus model its structured domains unambiguously (Figures 2A and 2B). As its human ortholog Rictor^19^, Avo3 is composed of three main domains (we retain Rictor nomenclature to facilitate their comparison): an N-terminal ARM domain (AD), a middle HEAT-like domain (HD) and the C-terminal domain (CD) (Figure 2B). The overall positioning of these domains in TORC2 is conserved with mTORC2 (Figure 2C). Both the AD and CD are located next to the Tor2 FKBP12-rapamycin binding (FRB) domain, with the AD interacting with the Avo1 N-terminal domain (ND) and part of the Tor2^KD^ (Figure 2A). Compared to Rictor, there is a large extension in the N-terminal region of Avo3 that is not resolved (residues 1-298), but it is predicted to contain two long alpha-helices connected to the AD by a linker (Figure S6A). However, this region is not essential for the function of TORC2^16^. The HD mostly interacts with Bit61, while also contacting the M-HEAT domain of the opposite Tor2 copy (Figure 2A). The HD is connected to the CD via a long linker, where Avo3 and Rictor structurally diverge. In Rictor, this linker includes a short, structured region that is positioned next to the Rictor^CD^, followed by the Zinc finger domain that precedes the CD (Figure 2C). These two features are absent in Avo3 (Figure 2B). In contrast, the Avo3 linker region next to the HD invades the FAT domain of Tor2 of the opposite monomer (Tor2’), and hence we name this region the Avo3 FAT-invading domain (FID; Figures 2A and 2B). The FID is connected to the CD by a disordered stretch of approximately 88 residues, giving it flexibility relative to both HD and CD. The insensitivity of TORC2 to rapamycin has been previously mapped to the Avo3^CD^, which, in agreement, precludes the binding of FKBP12-rapamycin to the Tor2^FRB^ (Figure 2D). Interestingly, the FRB is more occluded in mTORC2 due to the presence of the Rictor linker and part of mSin1 (“traverse”; see below), in addition to the Rictor^CD^ (Figure 2E).

**Figure 2.**
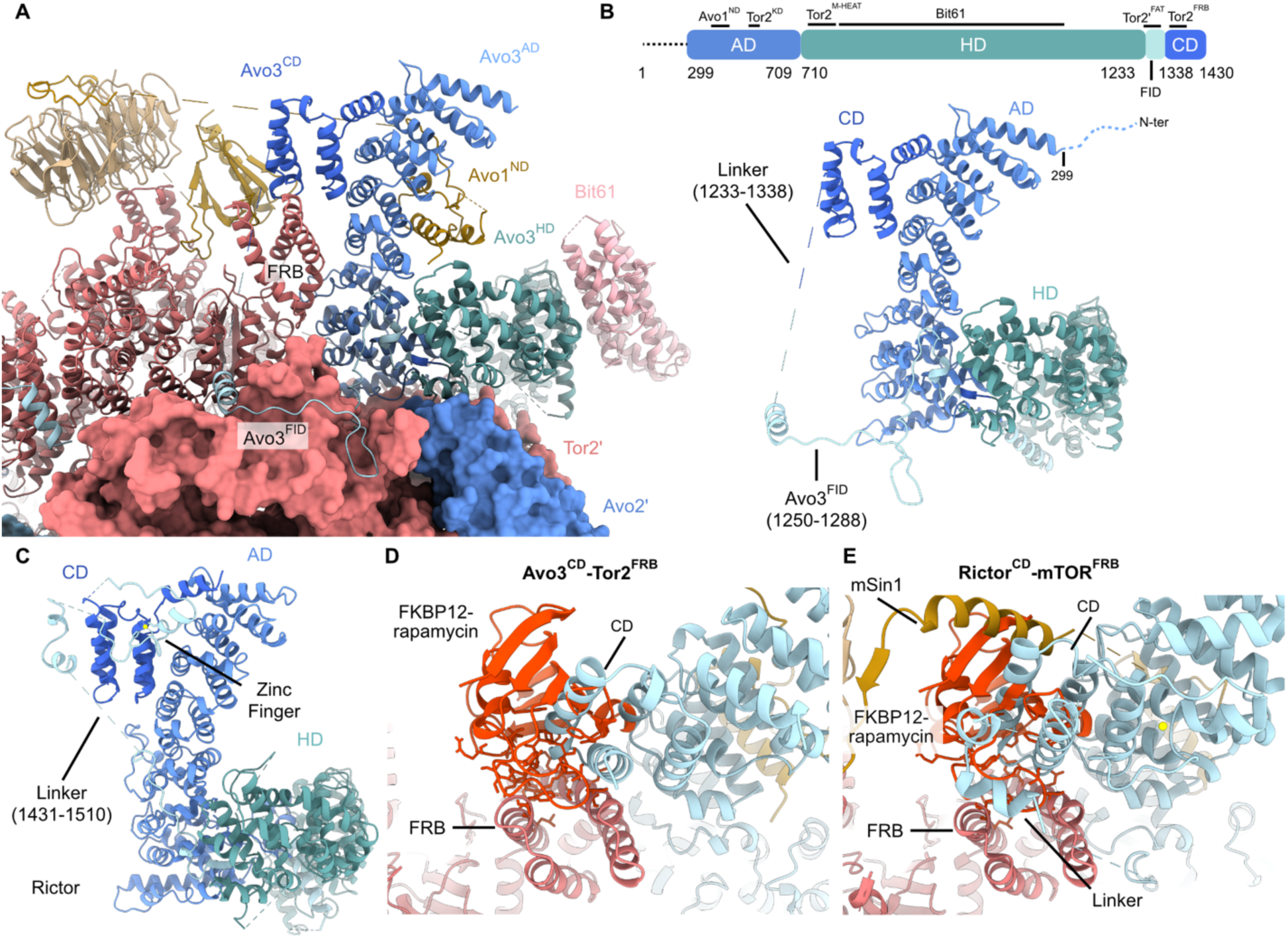
Structure of Avo3. **A.** Structure of Avo3 within TORC2. **B.** Model and schematic representation of Avo3 colored according to major domains. **C.** Model of Rictor (PDB: 6zwo) colored according to major domains. **D,E.** Overlay of FKBP12-Rapamycin complex (PDB: 4DRI) bound to the TOR FRB domain in TORC2 (**D**) and mTORC2 (**E**).

### Bit61 and Protor1 are orthologs

As inferred from the previous low-resolution TORC2 cryo-EM map^9^, Bit61 is located at the edges of the TORC2 dimer, interacting exclusively with the Avo3^HD^ (Figures 3A and 3B). Bit61 has a long unresolved N-terminal region (residues 1-287), followed by an HbrB domain that extends to the C-terminus (288-543). We have previously proposed that Bit61/Bit2 are orthologs of Protor1/Protor2^17^, but this has been contested due to the low sequence similarity^13^. To clarify this issue, we first compared our model to the AlphaFold prediction of Protor1. The predicted Protor1 structure shows high structural similarity to Bit61 (RMSD = 0.939 Å over 90 pruned atom pairs) (Figure 3C). We then tested whether AlphaFold could predict an interaction between Protor1 and Rictor. We included full-length Rictor in the prediction to reduce bias. In agreement with their structural conservation, the prediction shows that Protor1 binds exclusively to the Rictor^HD^ in a similar manner as Bit61 to the Avo3^HD^ (Figure S6B). Furthermore, the structural prediction could be readily fitted to a lower resolution mTORC2 cryo-EM map that contained Protor1^26^ in a similar place as in our reconstruction (Figure S6C,D). These observations are in line with previous biochemical data that show that Protor1 interacts specifically with Rictor^27^. Interestingly, only two residues in Avo3 and one in Bit61 that form hydrogen bonds are conserved with Rictor and Protor1, respectively (shown with an asterisk) (Figure 3B). Altogether, our data shows that despite the low sequence conservation, Bit61 and Protor1 are orthologous subunits.

**Figure 3.**
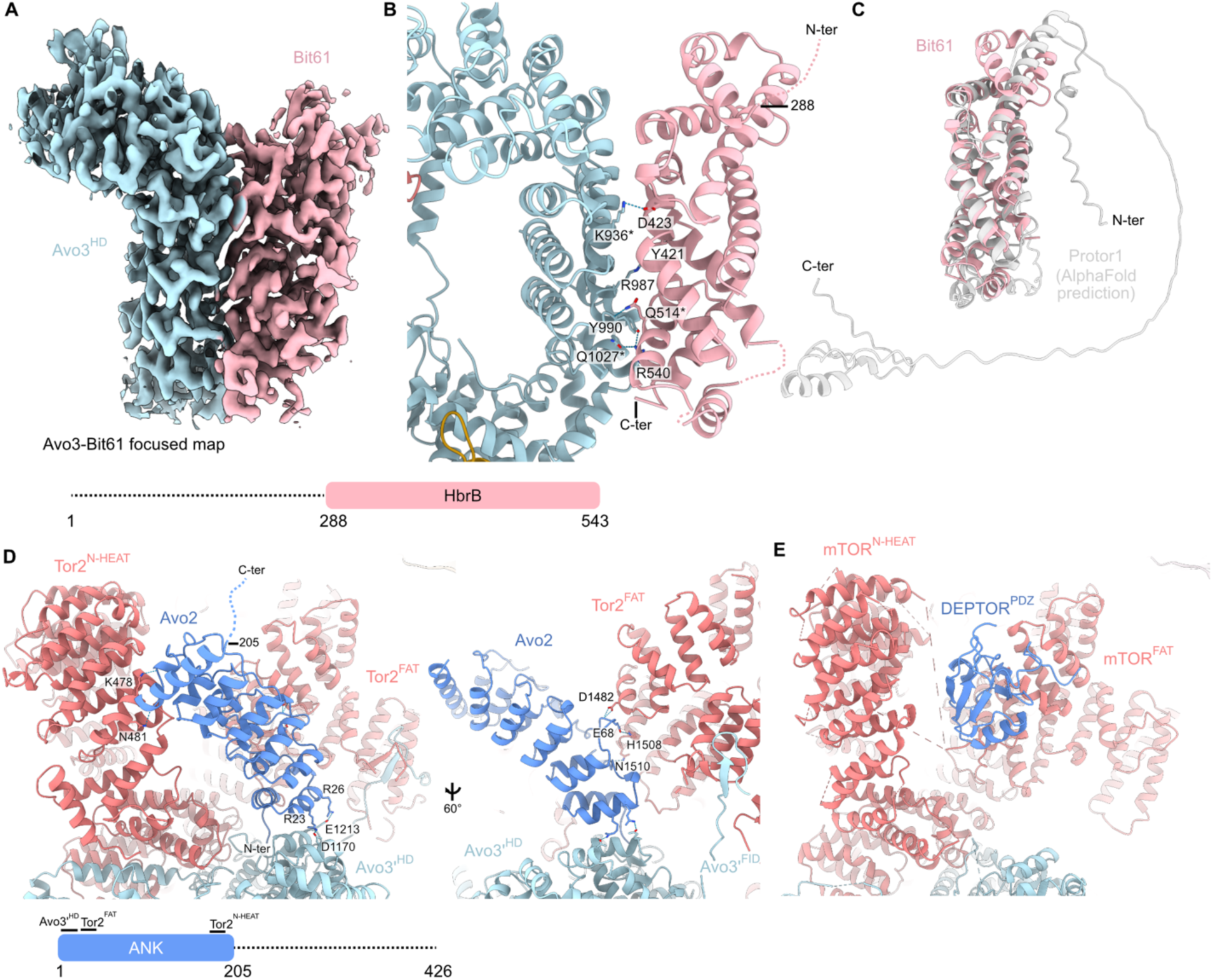
Structure of Bit61 and Avo2. **A.** Cryo-EM map of the TORC2 monomer focused on Avo3-Bit61. The schematic representation of Bit61 is shown below. **B.** Model of Bit61 bound to the Avo3 HD domain. Conserved residues with Rictor and Protor1 are indicated by an asterisk. **C.** Overlay between Bit61 and the AF3 prediction of Protor1. **D.** Model of Avo2. The schematic representation of Avo2 is shown below. **E.** Model of mTOR bound to DEPTOR (PDB: 7pe8). See also Figure S6.

### Avo2 interacts with the Tor2 kinase

In contrast to Bit61, which interacts exclusively with Avo3, Avo2 contacts both Tor2 and the neighboring Avo3 from the opposite monomer (Avo3’), effectively stabilizing the TORC2 dimer (Figure 3D). The structured N-terminal region of Avo2 is formed by ankyrin repeats (1-205; ANK), followed by a long, disordered stretch (206-426). The N-terminal end of Avo2 is inserted into the Avo3’^HD^, with two arginines from the second α-helix participating in hydrogen bonds with Avo3’ (Figure 3D). This region also interacts with the Tor2^FAT^, while the C-terminal part of the ANK domain interacts with the Tor2^N-HEAT^. In this manner, Avo2 bridges the N-HEAT and FAT domains of the Tor2 kinase. Interestingly, the position of Avo2 is reminiscent of where the PDZ domain of the mTORC1/2 regulatory protein DEPTOR binds^28^ (Figure 3E). However, the DEPTOR^PDZ^ only interacts with the mTOR^FAT^. The position of Avo2 is compatible with direct regulation of the catalytic activity of Tor2. Consistent with this idea, deletion of Avo2 renders yeast cells sensitive to myriocin, an inhibitor of sphingosine biosynthesis that induces compensatory TORC2 activation^18^. This defect is rescued by co-expression of a hyperactive allele of the main downstream TORC2 target, Ypk1 (Ypk1^D242A^), suggesting that Avo2 is necessary for TORC2 activation. Curiously, the isolated DEPTOR^PDZ^ has a weak stimulatory effect on mTORC1 activity *in vitro*^28,29^. Thus, it appears that both Avo2 and the DEPTOR^PDZ^, by interacting with the FAT domain of mTOR and Tor2, stabilize the kinase domain to promote the kinase activity of the complex.

### Avo1 domains are arranged around the TORC2 active site

Like mSin1, the N-terminal domain (ND, residues 1-70) of Avo1 binds to the Avo3^ND^, which stably anchors Avo1 to TORC2 (Figures S7A, 4A). In mSin1, an α-helical stretch termed the “traverse” bridges the active site and connects the ND to the Lst8 binding (LB) domain^19^ (Figure S7B). In Avo1, this region is much larger (553 amino acids) and is disordered (Figure 4A). Avo1 is predicted to have a helix that could correspond to the traverse, but it is prone to undergo a conformational change given its low pLDDT score (Figures S7C,D). The CRIM domain is flexibly linked to TORC2 via the LB (Figure 4A). Due to its high flexibility, the Avo1^CRIM^ is only resolved to low resolution, precluding unambiguous modeling. The Avo1^RBD^, which is adjacent to the CRIM, remains flexible and not visible. Finally, the C-terminal PH domain is located close to the Tor2 active site, interacting with both Tor2 and Lst8 (Figure 4A). The presence of the Avo1^PH^ in the active site is incompatible with ordering of the “traverse”, as it blocks the interface where mSin1 complements a β-sheet of mLst8.

**Figure 4.**
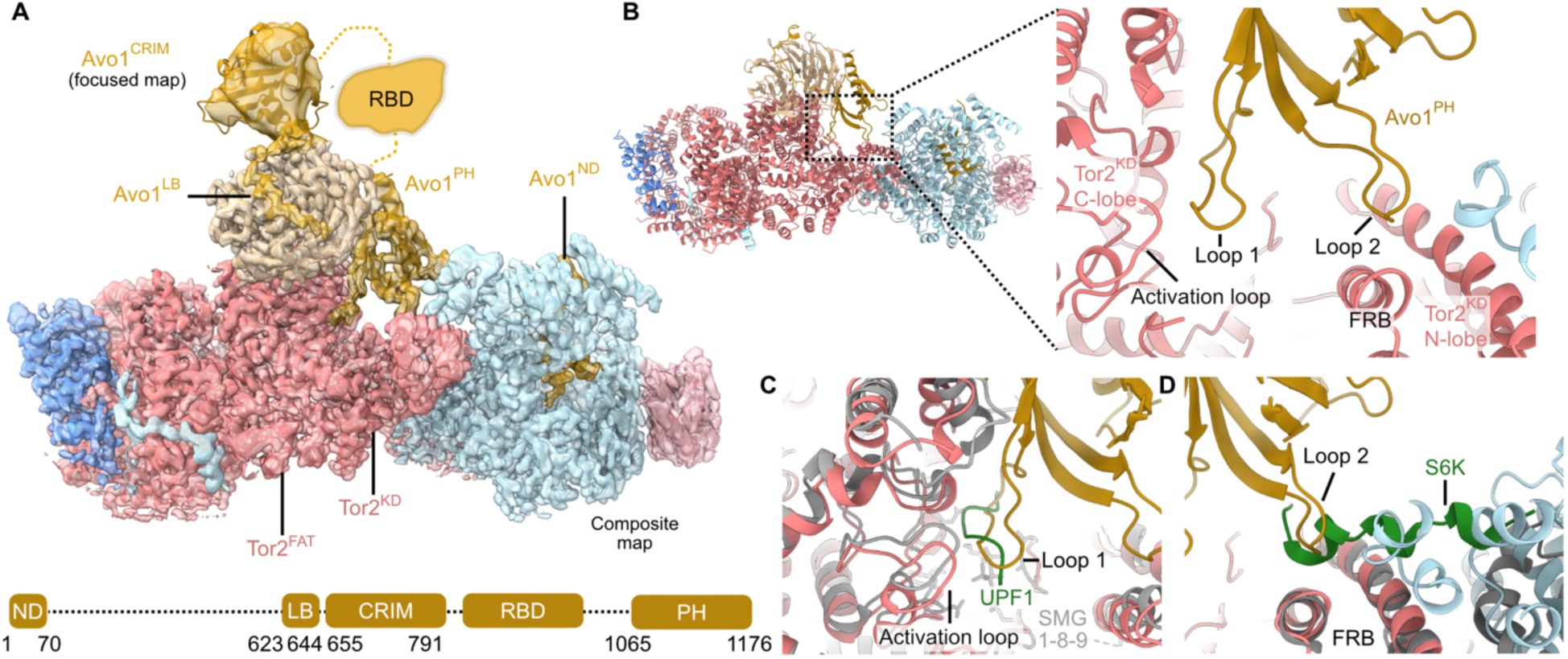
Structure of Avo1. **A.** Composite cryo-EM map of TORC2 containing the Avo1 PH domain in the active site cleft. The schematic representation of Avo1 is shown below. Low resolution density assigned to the Avo1 CRIM domain is overlayed. The CRIM domain is fitted best to the density, but the exact orientation remains uncertain due to extra disordered residues between the LB and CRIM not being observed. **B.** Zoom-in of the Avo1 PH domain inserted into the Tor2 kinase domain. **C.** Overlay between the Avo1^PH^-Tor2^KD^ and SMG1-8-9 bound to a peptide substrate, UPF1 (PDB: 6z3r) shows that Loop 1 would clash with the Tor2 substrate. **D.** Overlay between the Avo1^PH^-Tor2^KD^ and the structure of an S6K peptide bound to the mTOR FRB domain (PDB: 5wbh). See also Figure S7.

Within the catalytic cleft, Avo1^PH^ interacts with both lobes of the Tor2^KD^ using two loops. Loop 1 interacts with the Tor2 activation loop in the Tor2^KD^ C-lobe, while Loop 2 inserts into a pocket in the FRB in the Tor2^KD^ N-lobe (Figure 4B). Crucially, both interactions interfere with substrate binding to the active site. In the only structure of a phosphatidylinositol 3-kinase related kinase (PIKK; family of kinases that includes TOR) with bound substrate (SMG1-8-9 bound to UPF1^30^), the position of the substrate overlaps with Loop 1 (Figure 4C). Likewise, the binding of Loop 2 to the FRB overlaps with the position of an S6K peptide bound to mTOR^22^ (Figure 4D). Therefore, the observed position of the Avo1^PH^ is incompatible with substrate binding to Tor2. Given the structural similarity between the PH domains of Avo1 and mSin1 (Figure S7E), it is likely that similar interactions also occur in mTORC2, as previously proposed^14^, with rearrangement of the traverse being necessary.

### The Avo1 PH domain, and its binding to phosphoinositides, is required for TORC2 activation

Based on our structure and previous work^14^, we reasoned that deletion of the Avo1^PH^ would render TORC2 constitutively active and incapable of being downregulated by known inhibitory conditions. Surprisingly, this was not the case, as TORC2 in these cells appears to be hypomorphic, with cells presenting a lower steady state level of Ypk1 phosphorylation (Figure 5A) and sensitivity to myriocin (Figure 5B). Interestingly, while TORC2 inhibition by hyperosmotic shock^31^ or palmitoylcarnitine (PalmC)^3^ treatment was normal in cells lacking the Avo1^PH^ (Figures S8A and S8B), hyperactivation of TORC2 in response to hypoosmotic shock (Figure 5C) or myriocin treatment^32^ (Figure 5D), was no longer observed. Therefore, contrary to our prediction, the Avo1^PH^ appears to have a positive role on TORC2 activity. To better understand these results, we compared the structures of TORC2 classes with and without the Avo1^PH^ inserted into the active site. When bound to the Tor2^KD^, Avo1^PH^ pushes Lst8 away from the Tor2^FRB^, widening the active site cleft by ∼7 Å (Figures 5E and 5F). Further classification of particles lacking the Avo1^PH^ revealed that, in its absence, part of the Tor2 Negative Regulatory Domain (NRD), a long, disordered loop that has been shown to inhibit mTOR^33^, invades the active site and shields the activation loop (Figure S8C). Therefore, the Avo1^PH^ appears to prevent cleft closure and insertion of the NRD into the active site, both of which would restrict substrate access. This would explain the lower basal TORC2 activity in *avo1^ΔPH^* cells, their inability to be activated, and is consistent with previous data which shows that both the overexpression or deletion of the mSin1^PH^ reduces mTORC2 activity^14^.

**Figure 5.**
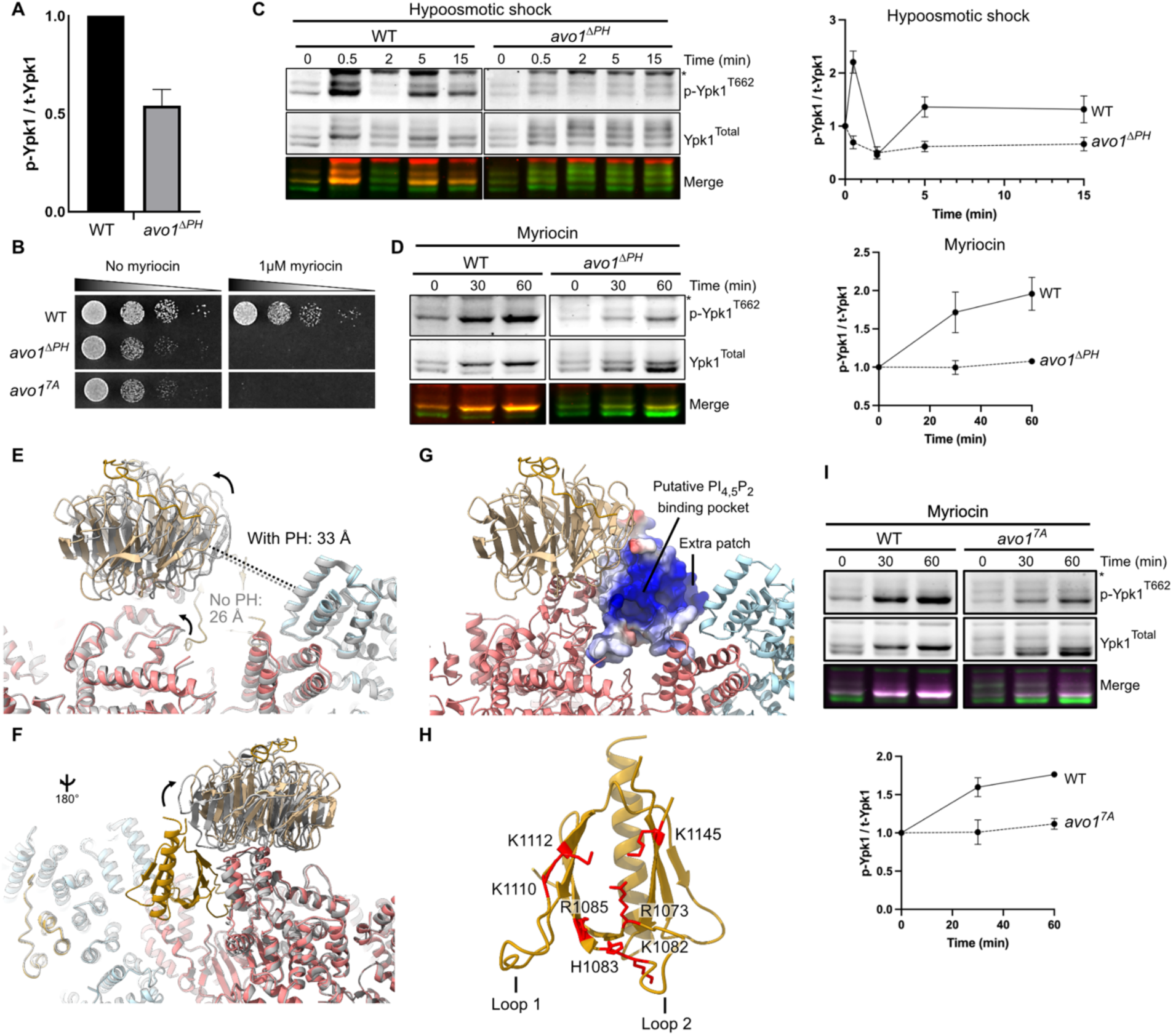
The Avo1 PH domain is required for TORC2 activation. **A.** Immunoblots of Ypk1 phosphorylation after activation of TORC2 by myriocin (5μM) in wild type and *avo1^ΔPH^*cells (n=3). Quantification is shown below. **B.** Immunoblots of Ypk1 phosphorylation after activation of TORC2 by hypoosmotic shock (1M to 0.1M sorbitol) in wild type and *avo1^ΔPH^*cells (n=3). Quantification is shown below. **C.** Quantification of basal TORC2 activity between wild type and *avo1^ΔPH^* cells (n=3). **D.** Growth assays of wild type, *avo1^ΔPH^* and *avo1^7A^* cells in the absence and presence of myriocin. **E,F.** Comparison of the conformation of the Tor2^KD^-Lst8 structure between TORC2 particles with and without (grey) the Avo1^PH^ in the active site. **G.** The putative canonical and non-canonical (extra patch) PI_4,5_P_2_ binding site of the Avo1^PH^, shown colored according to the electrostatic potential. **H.** Residues that form the PI_4,5_P_2_ binding site based on homology with other PH domains and that were mutated to alanines are indicated in red. **I.** Immunoblots of Ypk1 phosphorylation after activation of TORC2 by myriocin (5μM) in wild type and *avo1^7A^* cells (n=3). Quantification is shown below. Data are represented as mean ± SD. See also Figure S8.

For TORC2 to be activated, the Avo1^PH^ must be released from the catalytic cleft. It has been previously shown that the Avo1^PH^ is required for the regulation of TORC2 activity by PI_(4,5)_P_2_^16^. In our structure, the putative Avo1^PH^ PtdIns binding pocket^34^ is buried within the active site, inaccessible to PM-inserted PtdIns (Figure 5G). Of note, we identified an adjacent, potential “non-canonical” PtdIns binding site, also present in the PH domain of Slm1^35^ and Akt^36^ (“Extra patch” in Figure 5G). Binding of PtdIns to either site thus requires retraction of the Avo1^PH^ from the active site, which would provide a mechanism for TORC2 activation by PI_(4,5)_P_2_. To test this, we mutated the seven residues that form the positively charged PtdIns binding pocket (Figure 5H, *avo1^7A^*) and tested the response of these cells to myriocin treatment. In agreement with our hypothesis, like *avo1^ΔPH^* cells, *avo1^7A^* cells could not activate TORC2 (Figure 5I) and were sensitive to myriocin (Figure 5B). Therefore, as previously proposed for mTORC2^14^, our data suggest that PtdIns play a role in acute activation of TORC2 by removing the Avo1^PH^ from the active site. Curiously, this mechanism appears to not play a role in acute inactivation of TORC2 (c.f. Discussion).

### A positive pocket in Avo3 regulates TORC2 activity

The Avo1^PH^ is dispensable for TORC2 localization to the PM, and depletion of PI_(4,5)_P_2_, while causing loss of TORC2 activity, is not sufficient to remove TORC2 from the PM^16^. Consistently, both *avo1^ΔPH^* and *avo1^7A^* cells display normal TORC2 localization as discrete puncta on the PM (Figure 6A). Thus, regulation of TORC2 by PtdIns does not involve obvious changes in TORC2 recruitment to the PM, suggesting that binding of TORC2 to the membrane is mediated by a distinct interface of the complex. To identify potential membrane-binding regions, we analyzed the electrostatic potential and hydrophobicity of the TORC2 dimer. We identified a large positively charged patch (that we term “PM patch”) formed by the Avo3^HD^ and Tor2^M-HEAT^ domains, with additional contribution from the Tor2^N-HEAT^, that could mediate electrostatic interactions with the membrane (Figure 6B). Binding of TORC2 to the PM through this interface would be consistent with previous data showing an essential role of Avo3 for TORC2 localization at the PM^16^. Modeling of TORC2 binding to the PM using this surface results in an orientation reminiscent of mTORC1 positioning on the lysosomal membrane via the Rag GTPases (Figure S9A). Interestingly, a recent structure of membrane-bound mTORC1^37^ shows that Raptor (the mTORC1 counterpart of Avo3) interacts directly with the membrane using residues located in a similar position to the Avo3^HD^. We could not identify an obvious hydrophobic patch on the surface of TORC2 (Figure S9B), further arguing that membrane anchoring is mainly due to electrostatic interactions between positively charged residues in TORC2 and anionic lipids on the PM.

**Figure 6.**
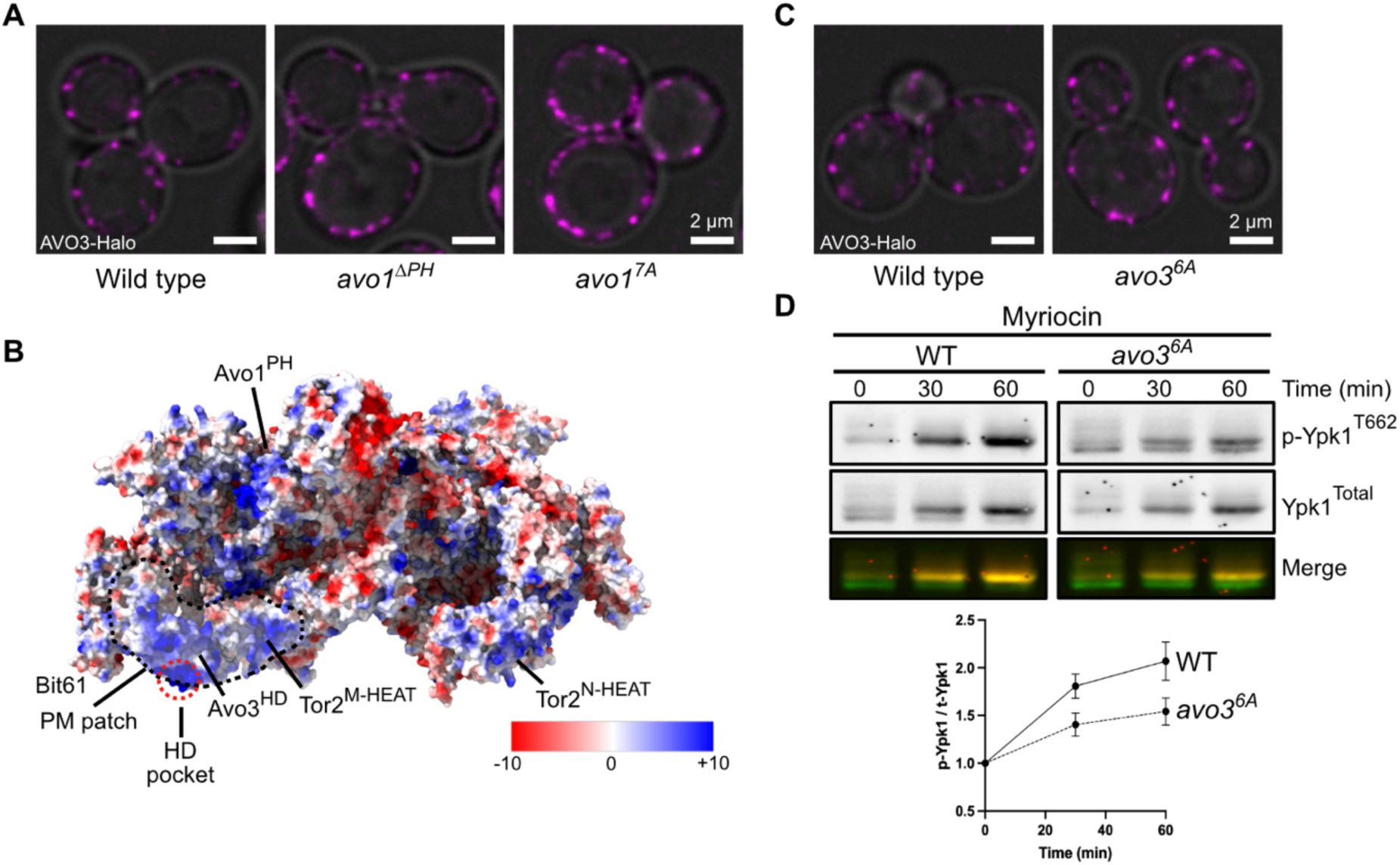
A positive pocket in Avo3 regulates TORC2 activity. **A.** Representative confocal microscopy images of Avo3-mScarlet-I3 in wild type, *avo1^ΔPH^* and *avo1^7A^* cells. Scale bar=2μm. **B.** Electrostatic potential map of the TORC2 dimer reveals a highly charged surface formed by Bit61, the Avo3^HD^ and the Tor2^M-HEAT^. **C.** Localization of TORC2 expressing *AVO3-HALO* in wild type and *avo3^6A^* cells. Scale bar=2μm. **D.** Immunoblots of Ypk1 phosphorylation after activation of TORC2 by myriocin (5μM) in wild type and *avo3^6A^* cells (n=3). Quantification is shown below. Data are represented as mean ± SD. See also Figure S9.

Within the PM patch, there is a highly charged pocket in the Avo3^HD^ that, in our model, would be positioned close to the membrane and accessible to lipids (Red circle in Figure 6B). Thus, we reasoned that this pocket (which we call “HD pocket”) might be particularly important for the regulation of TORC2 activity. However, likely due to the contribution of several regions to the PM patch, mutation of the HD pocket (alanine substitutions of 6 basic residues) alone was not sufficient to obviously affect TORC2 membrane binding (Figure 6C). Nevertheless, in these cells, TORC2 activation by myriocin was compromised to a similar extent as in *avo1^ΔPH^ and avo1^7A^* cells (Figure 6D). Thus, while membrane binding requires the contribution of different regions on the surface of TORC2, the HD pocket is important for TORC2 activation.

### Conformational changes of the Avo1 CRIM domain allow substrate delivery to the active site

Substrate engagement of TORC2 has been mapped to the CRIM domain of Avo1/Sin1^10^. We tentatively fitted the AlphaFold prediction of the Avo1^CRIM^ to our low-resolution density, guided by its position in the protein, just next to the Avo1^LB^ (Figures 4A and S10A,B). In this orientation, while not unambiguous, the Avo1^CRIM^ substrate-binding “acidic loop”^10^ faces away from Lst8 and is thus accessible for substrate recruitment (Figure 7A). The relative position of the substrate-binding loop would not be affected by errors in our fit, given that it is located opposite the Avo1^LB^-Avo1^CRIM^ junction. In the AlphaFold prediction of Lst8 with Avo1^CRIM^, the CRIM domain is rotated towards the Tor2^KD^ (Figure S10C) suggesting that the Avo1^LB^-Avo1^CRIM^ junction serves as a flexible hinge. Including Ypk1, Tor2 and Avo3 in the prediction showed an even larger structural transition in which the Avo1^CRIM^ binds to Ypk1 through an interface involving the “acidic loop”, and rotates inwards towards the active site, positioning the C-terminal region of Ypk1 in the catalytic cleft (Figures 7B and S10D). This conformation is consistent with known binding of Avo1/Sin1^CRIM^ to substrates^10^ and phosphorylation by TORC2 of residues in the C-terminus of Ypk1^5^. Together, our data and structural predictions suggest that the Avo1^CRIM^ can adopt distinct conformations to bind and present the substrate to the active site. This conformational transition is supported by reanalysis of previous cryo-EM maps of mTORC2 in the presence of its substrate Akt, in which the extra density assigned to the mSin1^CRIM^ undergoes similar changes as those observed between our cryo-EM map and the AlphaFold predictions (Figures S10E,F).

**Figure 7.**
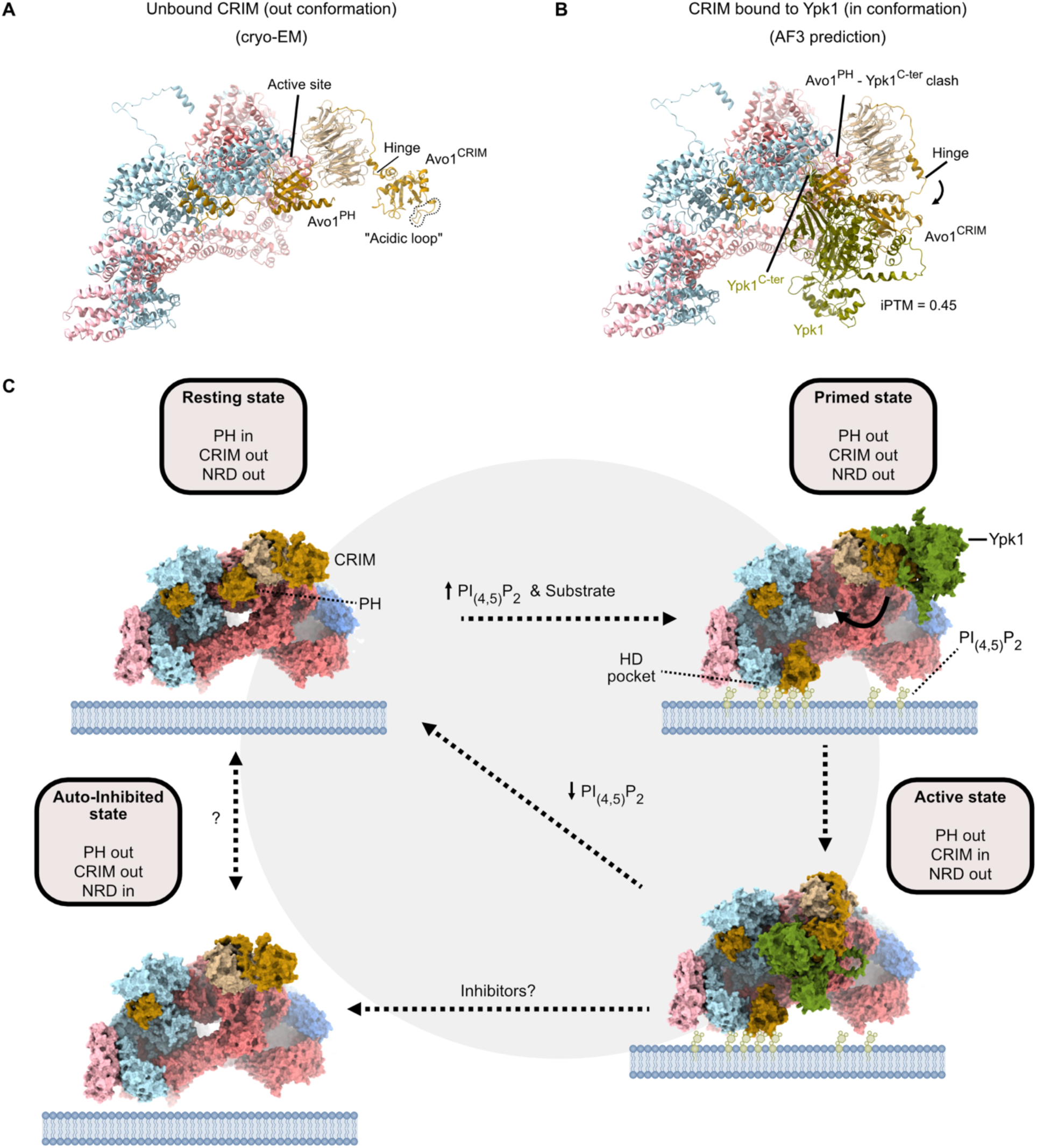
Proposed model of TORC2 activation. **A.** Position of the Avo1^CRIM^ based on the cryo-EM data on this study. **B.** AlphaFold3 prediction of Ypk1 bound to the Avo1^CRIM^, showing a rotation of the Avo1^CRIM^ towards Tor2, which would position Ypk1 in the active site. **C.** Model of TORC2 activation on the plasma membrane. Under low local levels of PI_4,5_P_2_ (resting state), positioning of the Avo1^PH^ in the Tor2 active site is favored (“in conformation”). Upon an increase in local PI_4,5_P_2_, binding of the Avo1^PH^ to PtdIns frees the active site (“out conformation”). In this state, the Avo1^CRIM^ can position Ypk1 in the active site, promoting its phosphorylation. TORC2 can also adopt an “inhibited state” in which the Tor2^NRD^ invades the active site. See also Figures S10 and S11.

## Discussion

### Model of TORC2 activation by Avo1 and Avo3

Collectively, we propose a model of TORC2 activation that requires the concerted action of the Avo1^PH^, the Avo1^CRIM^, Avo3 and PI_(4,5)_P_2_ (Figure 7C). Under basal conditions (“resting state”), positioning of the Avo1^PH^ near the active site (“in conformation”) is favored and the Avo1^CRIM^ points outwards, flexibly linked near the Tor2^KD^ and accessible for binding to Ypk1 or other substrates. When local levels of PI_(4,5)_P_2_ increase, for example, liberated by eisosomes upon increased membrane tension^38^, the Avo1^PH^ “out conformation” is favored and the active site becomes free (“primed state’”). Once the Avo1^PH^ is stabilized in its “out conformation” by PI_(4,5)_P_2_, the Avo1^CRIM^-bound substrate can rotate and deliver the substrate to the TORC2 active site (“active state”). Given that the absence of the Avo1^PH^ in the active site results in insertion of the Tor2^NRD^, we surmise that PI_(4,5)_P_2_-induced Avo1^PH^ withdrawal from the active site should be mechanistically coupled to substrate entry. Such a “coincidence detection” mechanism would prevent mis-activation of TORC2 in the absence of its substrate. The transition between “primed” to “active” states might be further promoted by the Avo3 HD pocket, which, through binding to PI_(4,5)_P_2_ or other lipids, may allosterically activate the kinase, as suggested for Raptor in mTORC1^37^. The HD pocket is conserved in Rictor, where it can bind ATP when incubated in a large excess, showing that it indeed can accommodate positively charged moieties^19^. The increase in local PI_(4,5)_P_2_ could also serve to co-localize the TORC2 activators Slm1/Slm2, which interact with PI_(4,5)_P_2_ via their PH domain, and with TORC2 via Bit61 and Avo2^39^. Consistently, our model of PM-bound TORC2 accommodates such interactions, as both Bit61 and Avo2 would be accessible to membrane-bound factors.

### Inhibition and activation of TORC2 are mechanistically uncoupled

While our work demonstrates a clear role of the Avo1^PH^ domain, and presumably PI_(4,5)_P_2_, in TORC2 hyperactivation (e.g. in response to hypo-osmotic shock), this domain appears not to be required for TORC2 inactivation (e.g. in response to hyperosmotic shock). Moreover, we previously^3^ observed that hyperosmotic shock triggers the release of Slm1/2 from eisosomes, but hyperosmotic shock did not trigger the reverse accumulation of Slm1/2 in eisosomes. These observations force us to conclude that distinct mechanisms exist to activate and inactivate TORC2. Indeed, in parallel work, we have discovered that TORC2 inhibition is coupled to an excess of free sterols in the PM^4^. These results were surprising to us as we did not expect that ostensibly similar (but opposite) stresses such as hypo- and hyper-osmotic shocks would provoke distinct signaling pathways as opposed to oppositely affecting a single molecular pathway.

Considering these observations, we were intrigued to observe a putative “auto-inhibited state” of TORC2 wherein the Tor2 Negative Regulatory Domain (NRD) occludes the activation loop (Figure 7C). We speculate that stabilization or adoption of this auto-inhibited state may contribute to TORC2 inhibition triggered by hyperosmotic shock, palmitoylcarnitine or other stresses^4^. Inhibition of kinase activity by a *cis*-acting regulatory domain, also known as the PIKK regulatory domain, is a common feature of PIKKs and well characterized through structural studies^40,41^. In contrast, the TOR^NRD^ has been observed only at low resolution in the *Km*TOR-Lst8 dimer structure (from the thermotolerant yeast *Kluyveromyces marxianus*), outside of the active site^42^. Nevertheless, the mTOR^NRD^ is necessary for the regulation of mTORC1 activity *in vivo*^33,43^. The mTOR^NRD^ contains several phosphorylation sites, one of which (Ser2448) is phosphorylated by Akt in response to growth factors^33^. However, there are no mapped phosphorylation sites in this region of Tor2 (Saccharomyces Genome Database, www.yeastgenome.org), which contains only a few serine and threonine residues. Thus, while it is possible that phosphorylation regulates the conformation of the Tor2^NRD^, our data rather suggests an important role of the Avo1^PH^ in the exclusion of the Tor2^NRD^ from the active site. Alternatively/additionally, TORC2 might be actively inhibited otherwise by factors that stabilize the insertion of the NRD into the active site. This would explain why TORC2 can still be inhibited in the absence of the Avo1^PH^. Unfortunately, we were unable to test our hypothesis as our attempts to remove the Tor2^NRD^ failed. The importance of this region in yeast TOR signaling remains to be determined.

### Functional interaction between Avo1 domains

In our cryo-EM data, we don’t observe a correlation between the positions of the Avo1^PH^ and the Avo1^CRIM^; given that these domains are separated by a long region that also includes the Avo1^RBD^, each domain can theoretically move independently. If and how the movement of both Avo1^PH^ and Avo1^CRIM^ is coordinated, potentially modulated by substrate binding, remains to be determined. In mTORC2, there appears to be a functional relationship between the mSin1^PH^ and mSin1^RBD^, with the mSin1^PH^ inhibiting the binding of the mSin1^RBD^ to Ras^11,12^. In yeast, Ras2 is proposed to promote the phosphorylation of Ypk1 by TORC2^13^, but the role of the Avo1^RBD^ has not been explored in detail. To gain insights into the functional relationship between Avo1 domains, we fitted the crystal structure of the mSin1^PH^-mSin1^RBD^-KRas complex^11^ to our model, using the PH domain as an anchor (Figure S10). This modeling shows that the Avo1^RBD^ would be able to interact with Ras1/2 only if the Avo1^CRIM^ is in the position observed in our cryo-EM maps; that is, away from the Tor2^KD^ (Figure S11A). Binding of PM-bound Ras2 to the Avo1^RBD^ might promote the retraction of the Avo1^PH^ from the active site and its interaction with PI_(4,5)_P_2_, allowing the movement of the Avo1^CRIM^ towards the active site, which would otherwise clash with Ras2 (Figure S11B). This model would explain the stimulatory effect of Ras2 on TORC2 activity, acting together with PI_(4,5)_P_2_. Nevertheless, this is speculative and should be experimentally tested.

## Conclusion

Our results suggest that upstream signals that regulate TORC2, such as increases in membrane tension, might do so by modulating PI_(4,5)_P_2_ enrichment around TORC2^38^, promoting the conformational transitions driving its activation. Whether additional interactions between the Avo1^CRIM^ or Ypk1/2 with regions around the Tor2^KD^ impact substrate delivery to the active site remains to be determined. Likewise, explaining the mechanism by which the Avo3 HD pocket and Ras1/2 activate TORC2 requires the structural elucidation of membrane- and Ras-bound TORC2.

## Resource availability

### Lead contact

Requests for further information and resources should be directed to and will be fulfilled by the lead contact, Robbie Loewith.

## Materials availability

This study did not generate new unique reagents. Strains used in this study are available from the lead contact upon reasonable request.

Maps and coordinates have been deposited in the Electron Microscopy Database (EMDB) and Protein Data Bank (PDB), with accession codes: TORC2 monomer with Avo1 PH (EMD-XXX/PDB:XXX), TORC2 dimer with Avo1 PH (EMD-XXX/PDB:XXX), autoinhibited TORC2 monomer (EMD-XXX/PDB:XXX), consensus TORC2 (EMD-XXX), TORC2 monomer focused (EMD-XXX), Avo3 focused (EMD-XXX), Bit61 focused (EMD-XXX), Lst8-Avo1 PH focused (EMD-XXX), Lst8-autoinhibited focused (EMD-XXX), Lst8-Avo1 CRIM focused map (EMD-XXX), Composite TORC2 monomer with Avo1 PH (EMD-XXX), Composite autoinhibited TORC2 (EMD-XXX).

## Acknowledgements

We thank the DCI Geneva (https://cryoem.unige.ch) and DCI Lausanne (https://dci-lausanne.ch/) for their support. R.L received support from the European Research Council (AdG TENDO), the Swiss National Science Foundation and the Canton of Geneva. L.T is supported by Grant PID2023-147101NA-I00 funded by MCIU/AEI /10.13039/501100011033 and by the European Union, ERDF “A way of making Europe”.

## Author contributions

R.L initiated the project. L.Z performed protein purification, cryo-EM grid preparation, and optimization. L.T processed and interpreted the cryo-EM data and built the models. M.G.T performed immunoblots, microscopy and cell growth experiments. A.B constructed yeast strains. L.T and R.L supervised all the work and prepared the manuscript with input from the other authors.

## Supplemental information

Document S1. Figures S1-S11 and Table S1.

## STAR Methods

### Yeast strains

Yeast strains used for this study are listed in Table S1. Point mutant strains were constructed using CRISPR-Cas9.

### TORC2 purification

TORC2 was purified from Δ*bit2* yeast cells carrying a TAP tag insertion in Bit61. Cells were grown in YPD medium until reaching an OD_600_ of 5-7, harvested by centrifugation at 6000 rpm for 10 minutes, and then flash frozen in liquid nitrogen and stored at −80°C. Cells were lysed and then resuspended in 1.5 volumes of extraction buffer (50mM PIPES pH 7, 300mM NaCl, 0.5mM CHAPS, 0.5mM DTT plus 1mM PMSF and 1X Complete protease inhibitor cocktail (-EDTA) (Roche)). The lysate was cleared by centrifugation at 12000 rpm for 10 minutes, and the supernatant was incubated with IgG-coupled Dynabeads M270 Epoxy (ThermoFisher Scientific) for 2 hrs at 4°C. Beads were washed 5 times with wash buffer (50mM PIPES pH 7, 300mM NaCl, 1mM CHAPS, 0.5mM DTT) at 4°C and incubated with TEV protease (0.1 mg/ml in elution buffer; 20 mM bicine pH8, 300 mM NaCl, 1 mM CHAPS, 0.5 mM DTT) for 1hr at 18°C. The eluate was collected at 4°C and used immediately for cryo-EM grid preparation.

### Cryo-EM grid preparation and data collection

Home-made graphene-oxide (GO) coated QuantiFoil Au 1.2/1.3 grids were used for sample preparation. GO (Sigma-Aldrich) was diluted with pure water to 0.2 mg/mL and the solution was spun for 6 seconds on a minicentrifuge. 3 µL of the cleared solution was incubated on glow discharged grids for 1 min, blotted, and washed with 20 µL of pure water three times (two times on the glow discharged side and one on the back). Excess water was removed with filter paper and grids were air dried for at least 1 hour. GO-coated grids were incubated for 3 minutes with 5 µL of freshly purified TORC2 in a Leica GP2 instrument set to 90% humidity and 4°C. The grid was blotted for 2 seconds with 2 mm angle offset.

cryo-EM data were acquired in a 300 kV Titan Krios equipped with a Falcon 4 direct electron detector and Selectris X energy filter (DCI Lausanne). Exposures were recorded with a total dose of 60 e/Å2 with 10 eV slit width, target defocus range from −0.6 to

−1.6 μm and a pixel size of 0.726 Å. A total of 16,725 exposures in EER format were collected.

### Cryo-EM data processing

Data processing was done by CryoSPARC 4.1.2^44^. Motion correction, CTF estimation, exposure curation and blob picking were done in CryoSPARC Live. A summary of the processing pipeline is shown in Extended Data Fig. 1. Picked particles were extracted with 800-pixel box size and binned two times for 2D classification. Good classes were selected and used as templates for template picking. After 2 rounds of 2D classification, good particles re-extracted with a box size of 640 pixels and resampled to 512 pixels and refined to obtain an initial 3D volume. This volume was then used to perform one round of heterogenous refinement in C1. The best class containing 705,208 particles was refined using non-uniform refinement and CTF refinement, followed by a 2D classification job to remove remaining bad classes. Selected 2D classes (616,122 particles) were subjected to reference-based motion correction, downsampled to 512-pixel box size and refined using non-uniform refinement to 2.4 Å resolution with C2 symmetry (consensus TORC2 dimer). These particles were symmetry-expanded in C2, and a mask encompassing one TORC2 “monomer” was used to locally refine particles to 2.32 Å resolution (monomer focused). These particles were then used to locally refine the monomer on Avo3, obtaining a reconstruction to 2.2 Å resolution (Avo3 focused), and on Avo3^HD^-Bit61 to 2.7 Å resolution (Bit61 focused).

To resolve the Avo1^PH^, initial good TORC2 particles (705,288 particles) were symmetry-expanded in C2 and subjected to focused 3D classification without alignment with 10 classes. The class with the best resolved Avo1^PH^ was locally refined, subjected to an additional round of focused 3D classification without alignment and the best class was refined again. To further improve the density, we removed duplicate particles, performed one round of 2D classification and selected the good classes to locally refine 51,729 particles to 3.07 Å resolution. We noticed that Lst8 remained very flexible in these particles, part of which interacts with the Avo1^PH^. We thus performed local refinement with a mask on Lst8, obtaining a 3.0 Å resolution reconstruction. Both cryo-EM maps were used to create a composite cryo-EM map. The same set of particles were refined in C2 to obtain the TORC2 dimer with PH domain visible in both active sites, although not resolved as good as in the monomer.

For the autoinhibited TORC2 conformation, we selected the only class from the 10 class-3D classification in which no density was observed near the active site (166,628 particles), even at low threshold. This class was locally refined, duplicate particles were removed, and particles were cleaned by one round of 2D classification. We finally obtained a 2.93 Å resolution reconstruction from 125,785 particles, with a focused refinement on Lst8 at 3.1 Å resolution. For visualization and interpretation of the part of the Tor2^NRD^ that becomes visible, we used the non-sharpened map, as sharpening eliminated clarity in this region due to the high flexibility. Likewise, overall refinement of the dimer in C2 blurred the density and was not clear in the opposite active site. For these reasons, we did not include a dimer model.

To better resolve the Avo1^CRIM^, monomer focused, symmetry-expanded particles were locally refined with a mask encompassing Tor2 and Lst8. Refined particles were sorted using focused 3D classification with a mask on Lst8 and an extra region above where the extra density was visible, iteratively by selecting the best class and increasing the resolution cut-off for classification. After 3 rounds of focused 3D classification, a class with 48,125 was refined to 3.83 Å resolution. This resolution reflects mostly Lst8 and surrounding parts of Tor2, as the Avo1^CRIM^ itself remained at lower resolution due to its flexibility.

All maps were sharpened automatically and using DeepEMhancer. We noticed that for the highest resolution maps, DeepEMhancer eliminated the high-resolution features, and therefore, we used the automatically sharpened maps. Figures were made using UCSF ChimeraX 1.8.

### Model building

Structure predictions of all TORC2 components (Tor2, Lst8, Avo1, Avo2, Avo3, and Bit61) were downloaded from the AlphaFold Protein Structure Database (https://alphafold.ebi.ac.uk/) and used as initial models. Models were cut into rigid domains and fitted into the density in UCSF Chimera 1.16. The model was then manually built in Coot 0.8.9. Real space refinement was done in Phenix 1.19.2-4158. We used each focused map to first refine the corresponding region of the complex. Then, models were real space refined into the monomer focused map for both PH and “autoinhibited” TORC2 maps. Finally, to create the TORC2 dimer, the monomer was rigid body fitted into the opposite copy of the complex and real space refined against the full map.

### Fluorescence microscopy

To localize TORC2 in cells, Avo3 was endogenously tagged with a Halo tag. WT or mutant yeast cells were grown to mid log phase in low fluorescence media. For Halo-tag labelling with JF646, the liquid culture was supplemented with 200 nM Halo-tag ligand (#GA1121, Promega, 200 µM stock in DMSO) and incubated at 30°C for 30-120 min. Before imaging, cells were pelleted by gentle centrifugation, resuspended in fresh media without ligand, and mounted onto Concanavalin A-coated VI 0.4 µ-Slides (Ibidi) primed with media. Images were recorded at room temperature on a Leica STELLARIS 8 FALCON FLIM Microscope with a 63×1.4 Oil immersion objective and LAS X (Ver. 4.6.1.27508) software, and subsequently deconvolved with Lightning deconvolution (Leica). Images were processed using FIJI (ImageJ 1.54p).

### Spot assays

Saturated overnight yeast cultures (30 °C, SC medium) were diluted to an OD_600_ of 0.1 in the morning and grown into mid log phase (OD_600_ = 0.5–0.8). Log phase cells were diluted to an OD_600_ of 0.1, and a tenfold dilution series was spotted onto SC medium plates containing 1 µM myriocin (Sigma M1177, 2.5 mM in MeOH), or vehicle. Plates were incubated at 30 °C, and imaged when differences were most apparent (typically after 48 h for control, 72 h for myriocin).

### Western blot

Yeast cells were grown at 30°C in synthetic media (SC) buffered at pH 6.2 with 0.1 M Sorensen Buffer and supplemented with 2% glucose and appropriate amino acids and nucleobases. All experiments were performed in logarithmically growing cells (same OD_600_ between different strains).

For osmotic shocks, yeast cells were either grown in regular media and then diluted with prewarmed media containing 2M sorbitol to a final concentration of 1M sorbitol (hyperosmotic shock) or grown in media containing 1M sorbitol and then diluted with prewarmed media without sorbitol for a final concentration of 0.1M sorbitol (hypoosmotic shock). For specific treatments, cultures were supplemented either with 5 µM myriocin (Sigma M1177, 2.5 mM in MeOH), or 5 µM palmitoylcarnitine (Sigma P4509, 10 mM in DMSO). Samples were harvested at the indicated timepoints and processed according to standard TCA-urea extraction procedures. Protein lysates were separated on 7.5% or 10% SDS-page gels and blotted onto nitrocellulose membranes using the iBlot2 Gel Transfer System (Thermo Fisher Scientific). Membranes were blocked with BSA, incubated with primary antibodies (polyclonal goat anti-Ypk1 (Agrobio homemade; 1:25′000); monoclonal mouse anti-phospho-Ypk1^T662^ (homemade; 1:500), overnight at 4 °C, washed and incubated with secondary antibodies for 1 hour at room temperature, using PBS- or TBS-based buffers. Membranes were developed using an Odyssey imaging system (LI-COR) and signal intensities were quantified using FIJI (ImageJ 1.54f). Calculations were performed in Microsoft Excel and plotted using GraphPad.

