## Supplemental information for "Structural basis for TORC2 activation"

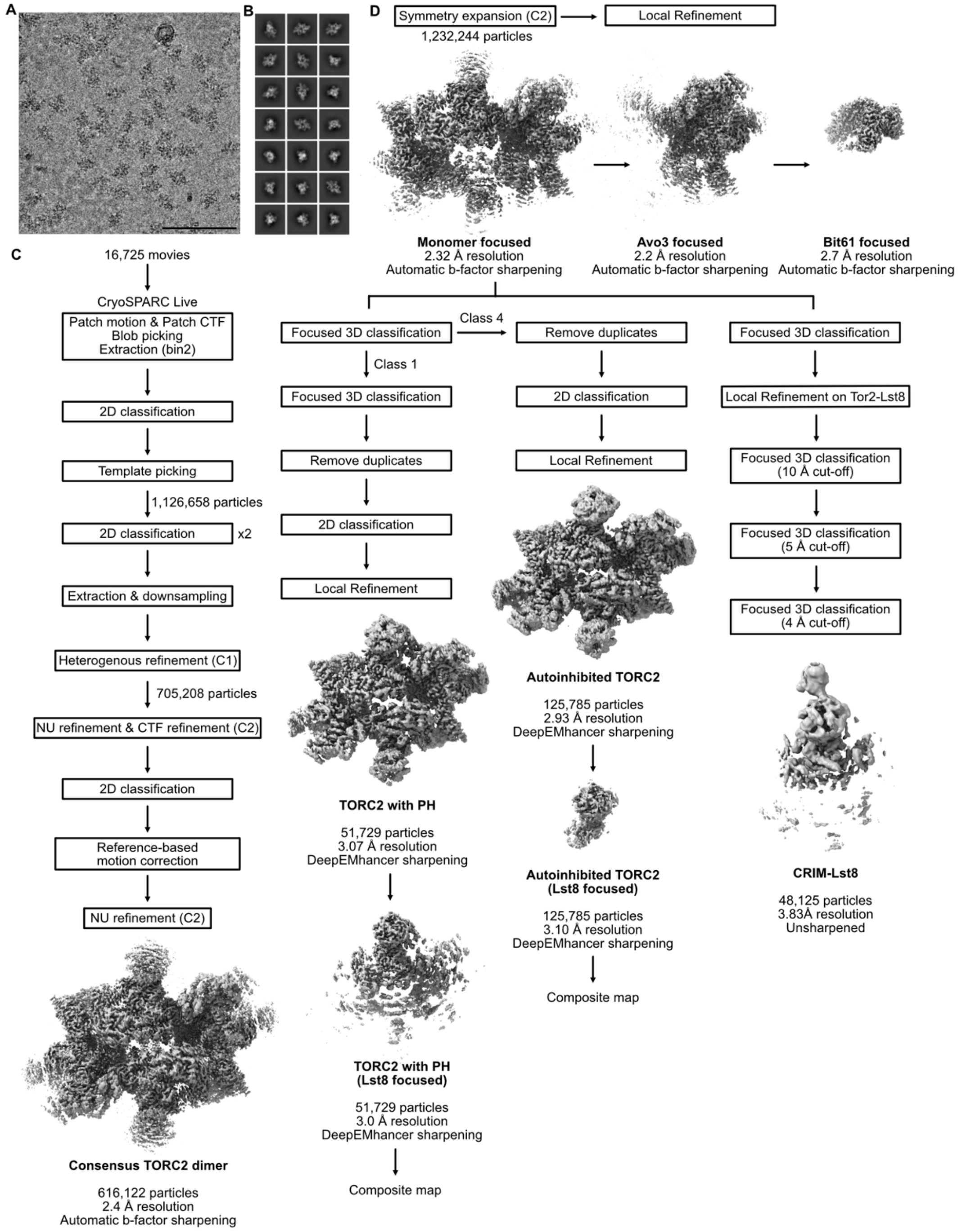


**Supplemental Figure 1. Cryo-EM data processing. A.** Representative cryo-EM micrograph. Scale bar = 100 nm. **B.** Representative 2D class averages. **C.** Processing pipeline of cryo-EM movies to obtain the consensus TORC2 dimer. **D.** Processing pipeline of cryo-EM movies to obtain the different focused maps.

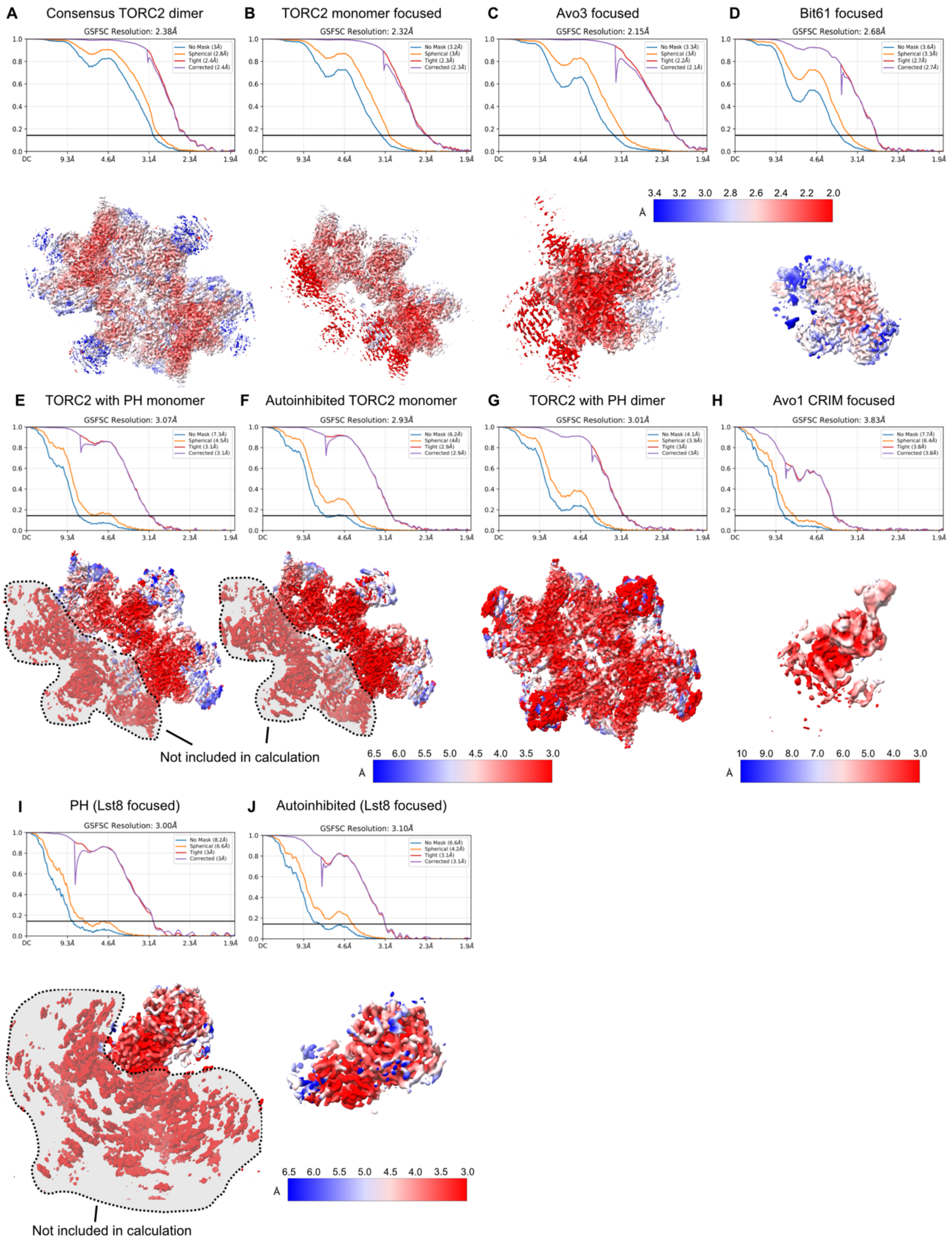


**Supplemental Figure 2. Cryo-EM validation and local resolution. A.** Half-map FSC and local resolution estimate for the consensus TORC2 dimer. **B.** Half-map FSC and local resolution estimate for the TORC2 monomer map. **C.** Half-map FSC and local resolution estimate for the Avo3 focused map. **D.** Half-map FSC and local resolution estimate for the Bit61 focused map. **E.** Half-map FSC and local resolution estimate for the TORC2 with PH monomer. **F.** Half-map FSC and local resolution estimate for the autoinhibited TORC2 monomer. **G.** Half-map FSC and local resolution estimate for the TORC2 with PH dimer. **H.** Half-map FSC and local resolution estimate for the Avo1 CRIM focused map. **I.** Half-map FSC and local resolution estimate for the TORC2 monomer with PH-Lst8 focused map. **J.** Half-map FSC and local resolution estimate for the autoinhibited TORC2 monomer-Lst8 focused map.

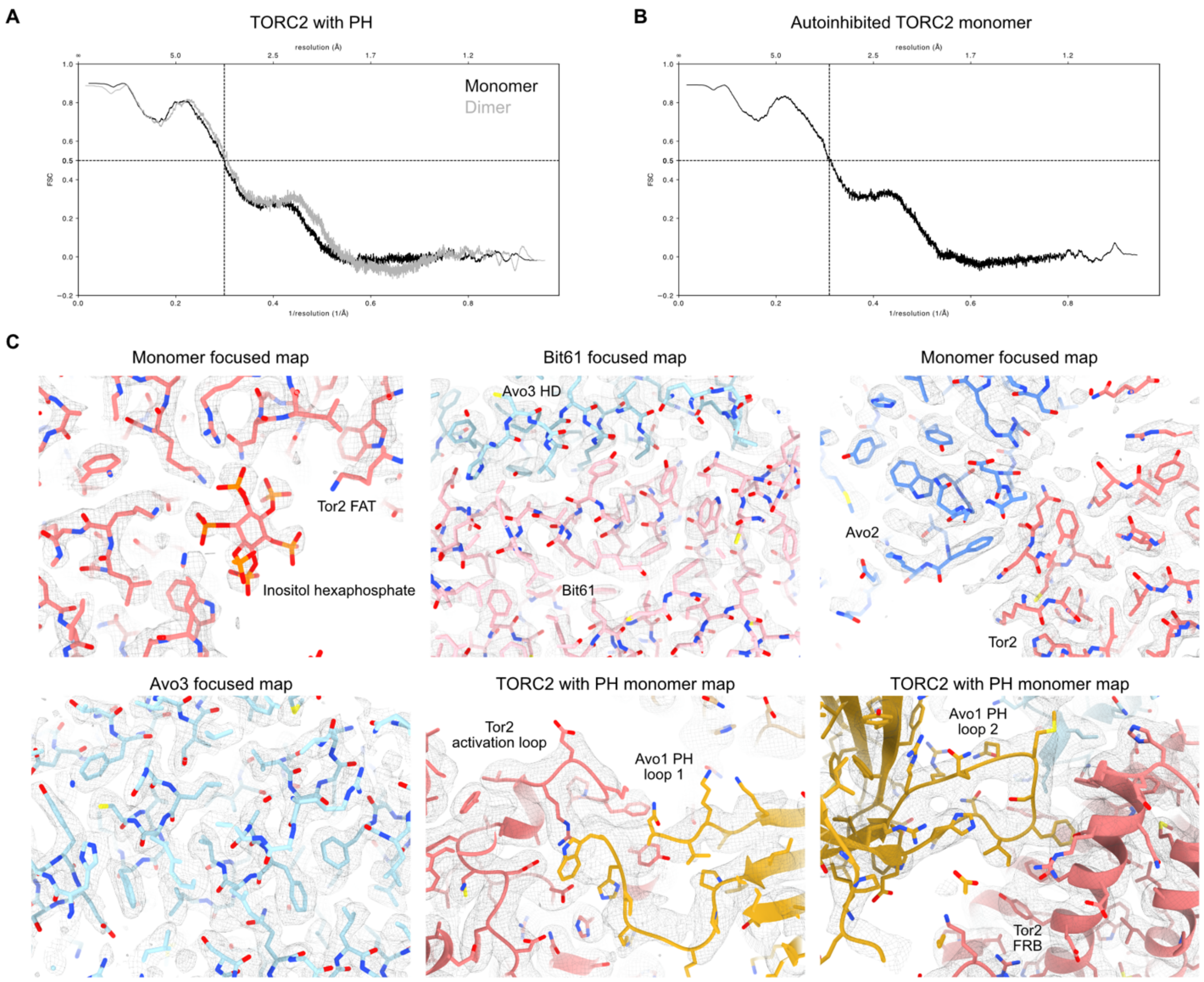


**Supplemental Figure 3. Model validation. A.** Model-map FSC curves for the TORC2 with PH domain monomer (black) and dimer (grey). **B**. Model-map FSC curves for the autoinhibited TORC2 monomer. **C.** Representative cryo-EM densities of different parts of TORC2.

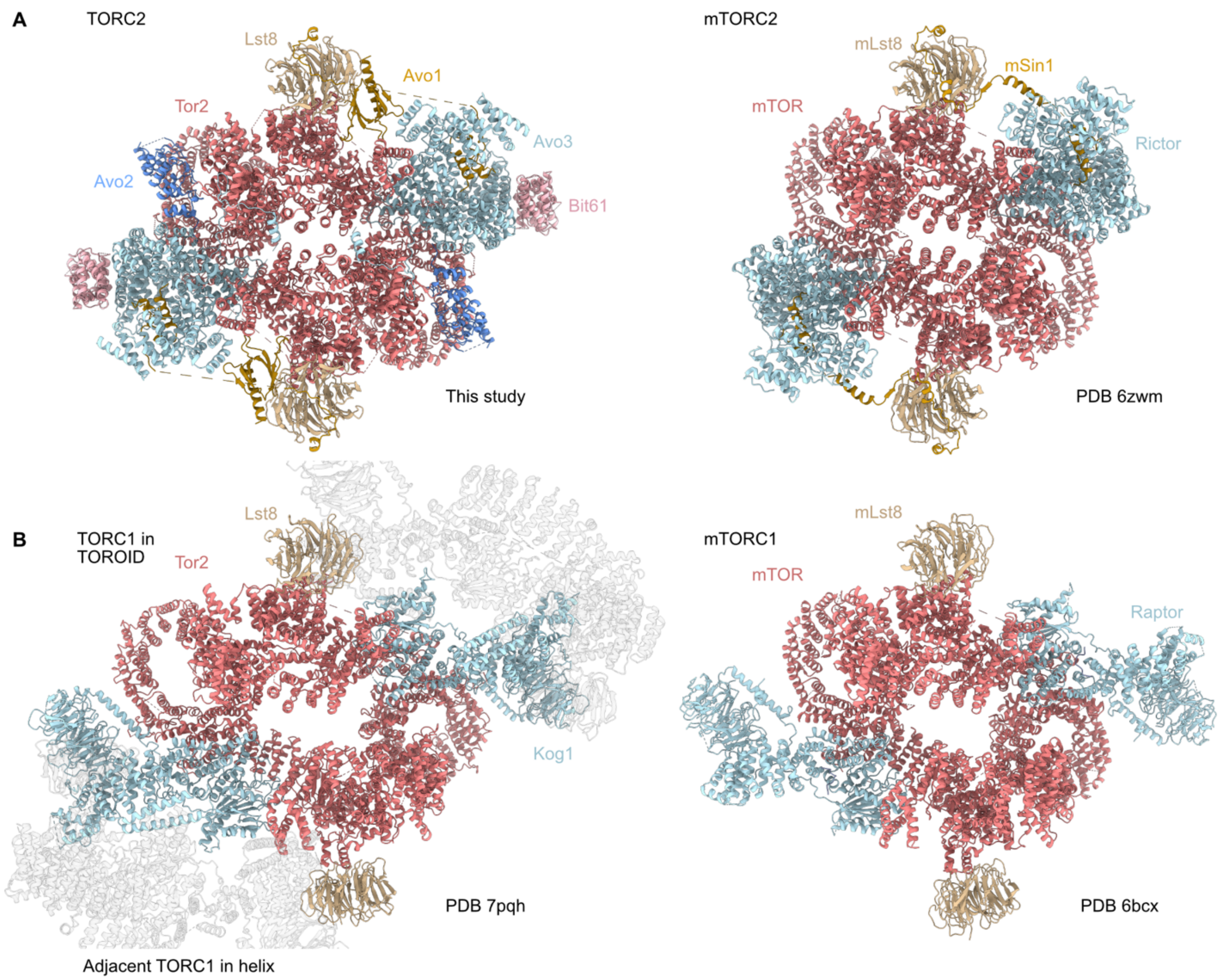


**Supplemental Figure 4. Comparison of (m)TORC1 and (m)TORC2. A.** Comparison of the structure of TORC2 (left, this study) and mTORC2 (right, PDB: 6zwm). **B.** Comparison of the structure of TORC1 (left, PDB: 7pqh) and mTORC1 (right, PDB: 6bcx). All models were aligned on Tor2 for their comparison.

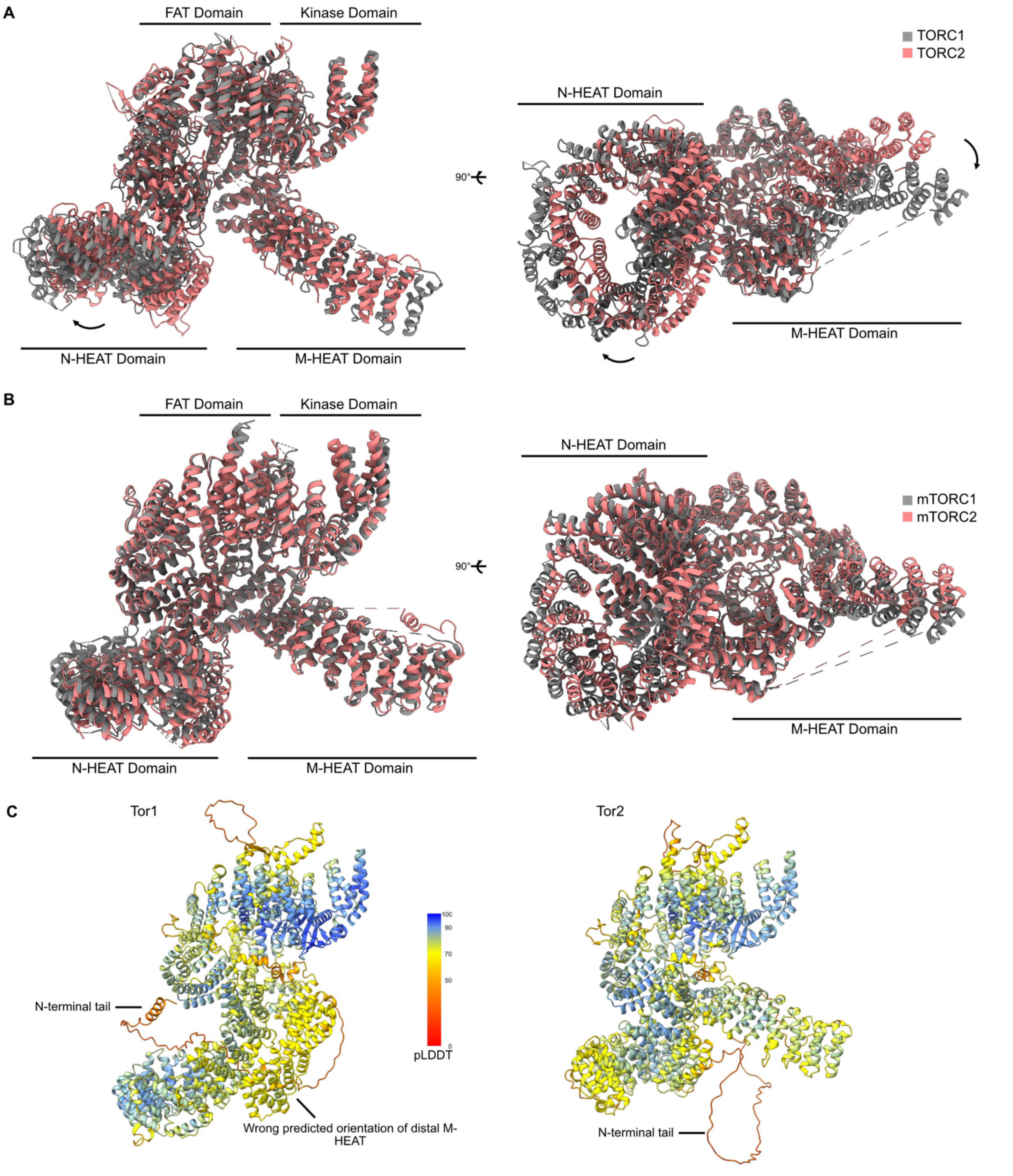


**Supplemental Figure 5. Structure of TOR and mTOR in (m)TORC1 and (m)TORC2.** **A.** Comparison of Tor2 when assembled in TORC1 (grey) or TORC2 (light coral). **B.** Comparison of mTOR when assembled in mTORC1 (grey, PDB: 6bcx) or TORC2 (light coral, PDB: 6zwm). **C.** AlphaFold predictions of Tor1 and Tor2, colored according to the pLDDT score. The position of the M-HEAT in the Tor1 prediction is incompatible with its incorporation into TORC1. Both also display a disordered N-terminal tail.

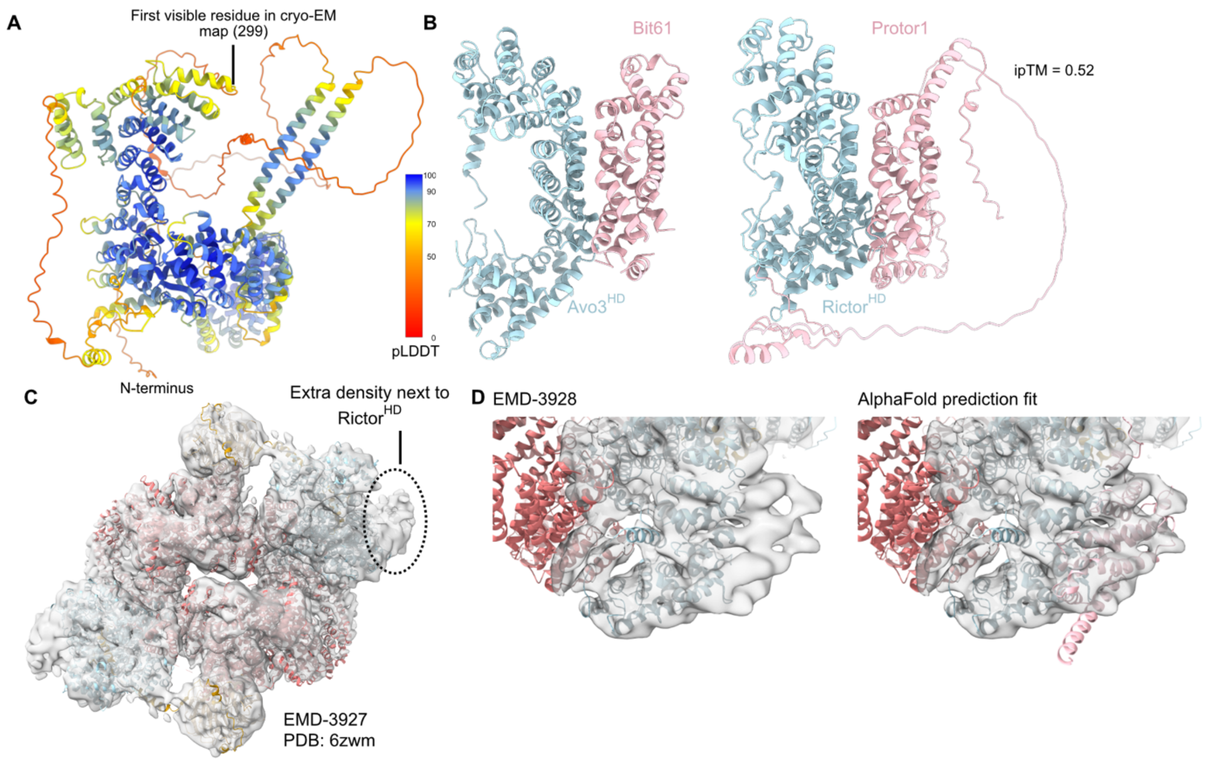


**Supplemental Figure 6. Protor1 is a Bit61 ortholog. A.** AlphaFold2 prediction of Avo3, showing the flexible N-terminal extension that is not visible in the cryo-EM map. **B.** Comparison between the interaction between Avo3 and Bit61, and the predicted interaction between Rictor and Protor1. **C.** Cryo-EM map of mTORC2 containing Protor1 (EMD-3927) fitted with the mTORC2 model (PDB: 6zwm). Extra unassigned density is located next to the Rictor$^{HD}$. **D.** Structural alignment of the Rictor$^{HD}$ between the TORC2 model and the Rictor-Protor1 prediction fits the unassigned density with Protor1, despite the low resolution (EMD-3928).

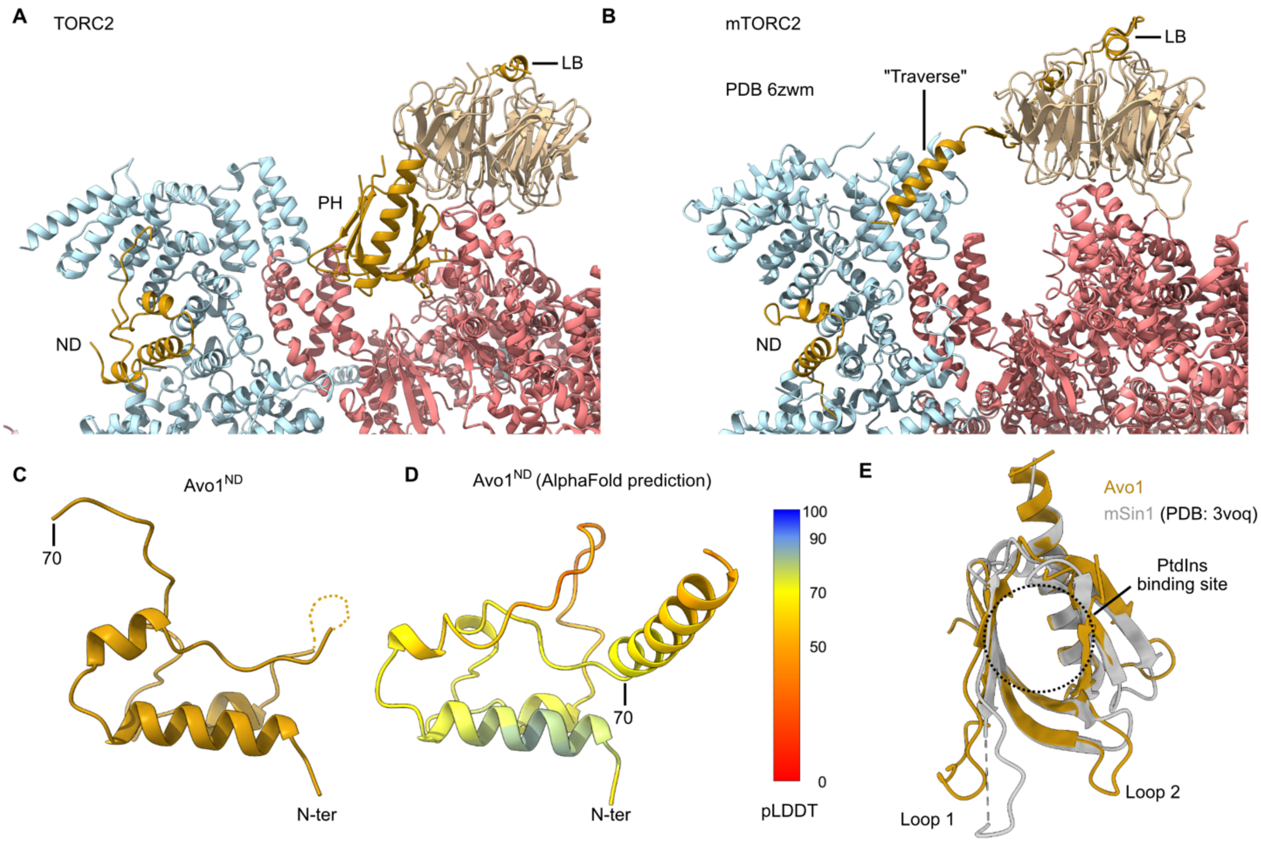


**Supplemental Figure 7. Comparison between Avo1 and mSin1. A.** Model of TORC2 (this study), showing the positions of the ND, LB and PH, with an absent "Traverse". **B.** Model of mTORC2 (PDB: 6zwm) showing the mSin1 "Traverse", which spans the active site cleft and links the ND with the LB. **C, D.** Model of the Avo1$^{ND}$ in the cryo-EM structure (**C**) and the AlphaFold prediction (**D**). The AF prediction suggests the presence of a helix that could be akin to the mSin1 "Traverse" after the last resolved residue in our structure. **E.** Structure alignment of the PH domain of Avo1 and mSin1 (PDB: 3voq), showing the putative PtdIns binding site.

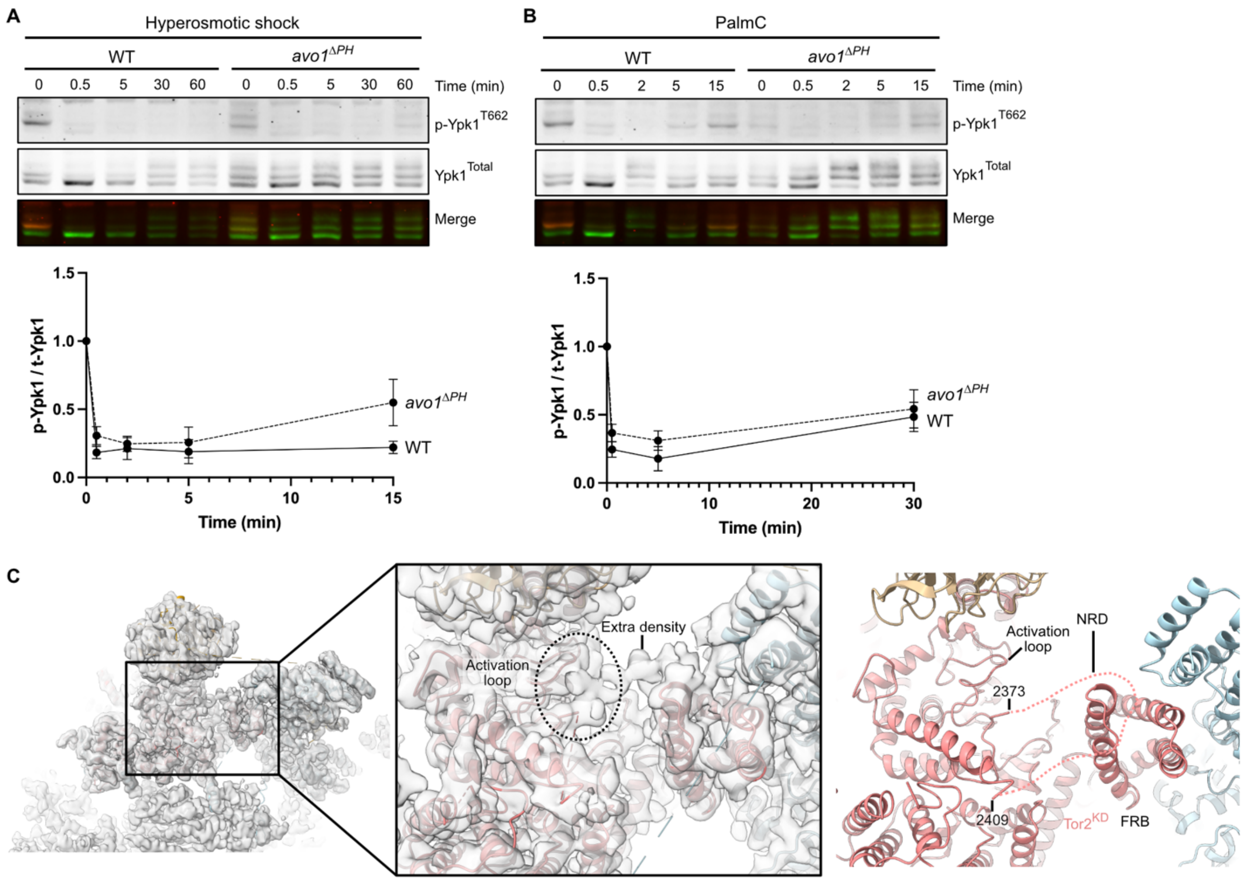


**Supplemental Figure 8. The Avo1 PH domain is not essential for acute inhibition of TORC2.** **A.** Immunoblots of Ypk1 phosphorylation after inhibition of TORC2 by hyperosmotic shock (0 to 1M sorbitol) in wild type and *avo1*$^{\Delta PH}$ cells (n=3). Quantification is shown below. **B.** Immunoblots of Ypk1 phosphorylation after inhibition of TORC2 by palmitoylcarnitine (PalmC) treatment (5μM) in wild type and *avo1*$^{\Delta PH}$ cells (n=3). Quantification is shown below. **C.** Zoom-in of an extra density, that could correspond to the Tor2 Negative Regulatory Domain (NRD), shielding the activation loop and extending towards the Tor2$^{FRB}$. Data are represented as mean ± SD.

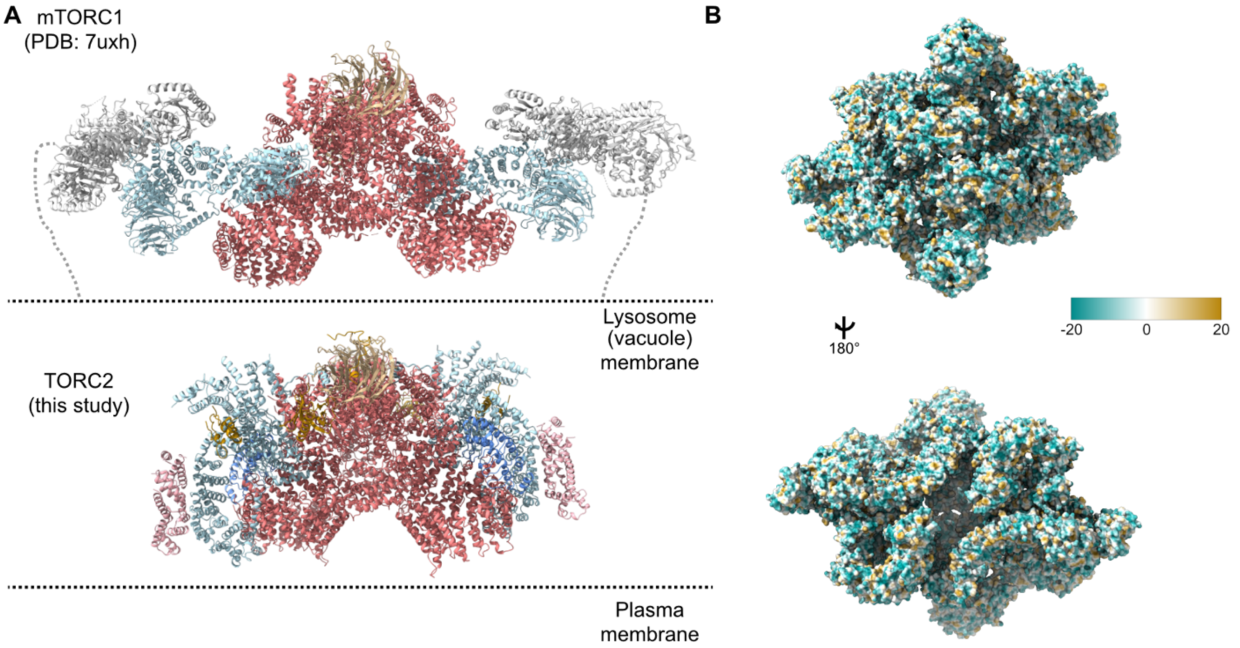


**Supplemental Figure 9. TORC2 membrane binding. A.** Comparison of membrane binding between mTORC1 (based on PDB: 7uxh) and TORC2. The orientation of TORC2 relative to the membrane is based on the positively charged surface identified on the complex. **B.** Hydrophobicity of TORC2 does not show any obvious region that could contribute to membrane binding. Higher values reflect higher hydrophobicity.

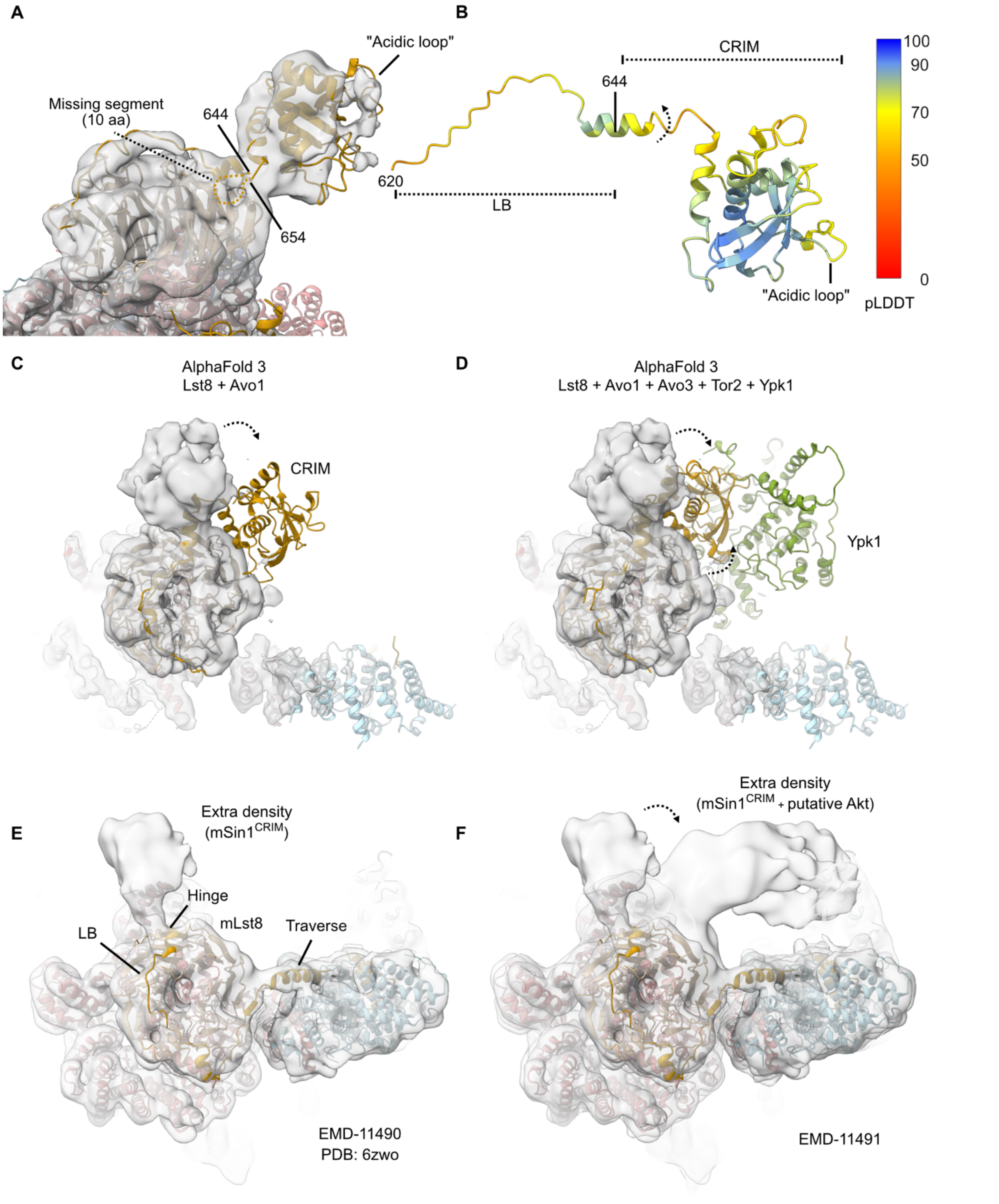


**Supplemental Figure 10. Fit and conformational flexibility of the Avo1 CRIM domain. A.** Cryo-EM density of the Avo1$^{CRIM}$ in the Avo1$^{CRIM}$-focused map, showing the last assigned residue in the Avo1$^{LB}$. **B.** AlphaFold2 prediction of the Avo1$^{LB}$ and Avo1$^{CRIM}$

domains showing the rotation hinge of the CRIM relative to the LB observed in our map. The fitted position is the best based on the cryo-EM density; however, approximately 10 to 15 residues between the start of the folded CRIM domain and the last visible residue are unaccounted for and should fold for the CRIM to be in the modeled position. **C.** AlphaFold3 prediction of the $Avo1^{CRIM}$ when predicted only with Lst8. **D.** AlphaFold3 prediction of the $Avo1^{CRIM}$ when predicted in the context of Lst8, Avo3, Tor2 and Ypk1. **E.** Extra density near Lst8 visible in mTORC2 (EMD-11490). **F.** A larger extra density is observed in the presence of Akt (EMD-11491), with movement of the extra density similar to the transition observed between our cryo-EM map and the AlphaFold3 prediction.

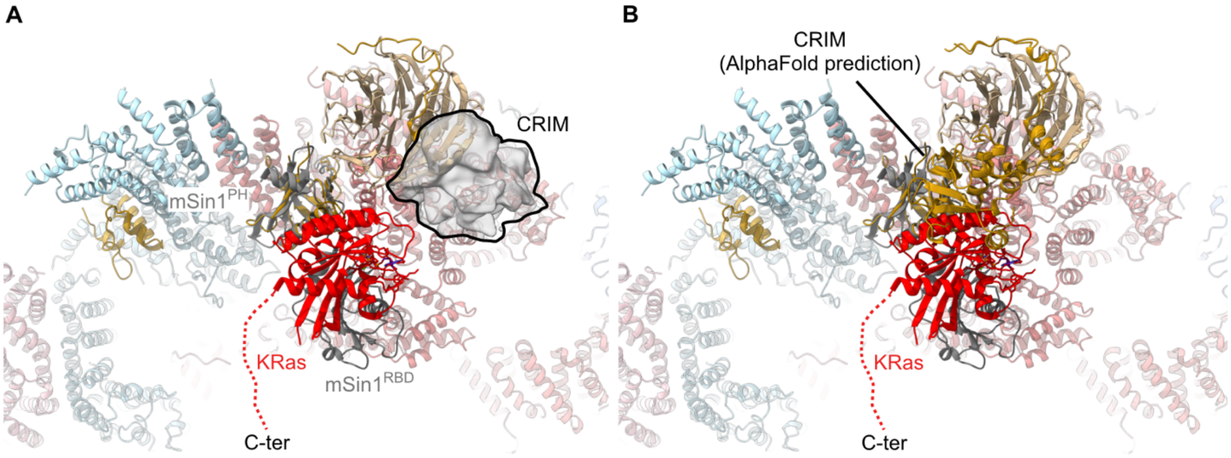


**Supplemental Figure 11. Modeling of Ras binding to TORC2. A.** Fit of the mSin1$^{RBD}$-mSin1$^{PH}$-KRas crystal structure (PDB: 7lc1) onto our model, aligned on the Avo1$^{PH}$. **B.** The modeled position of KRas clashes with the predicted position of the CRIM domain.

| Strain | Genotype | Source |
|---|---|---|
| TB50a | *a leu2 ura3 rme1 trp1 his3Δ GAL+ HMLa* | Loewith Lab |
| TB50α | *α leu2 ura3 rme1 trp1 his3Δ GAL+ HMLa* | Loewith Lab |
| CG1-34 | TB50α *bit2::kanMX6 Bit61-TAP::TRP1* | Gaubitz et al. 2015[1] |
| MTY028 | TB50a *avo1-ΔPH::T(ADH1)-KanMX6* | This study |
| #4663 | TB50a *avo1-R1073A/ L1082A/H1083A/R1085A/K1110A/K1112A/K1145A AVO3-Halo::KanMX* | This study |
| #4484 | TB50α *avo3-K784A/ R787A/R788A/R794A/ R797A/R799A* | This study |
| #3294 | TB50a *AVO3-Halo::KanMX* | This study |
| MGTY058 | TB50α *avo1-ΔPH:: (ADH1)-HphMX AVO3-Halo::KanMX* | This study |
| MGTY109 | TB50a *avo1-R1073A/ L1082A/H1083A/R1085A/K1110A/K1112A/K1145A AVO3-Halo::KanMX* | This study |
| #4488 | TB50a *avo3-K784A/ R787A/R788A/R794A/ R797A/R799A-Halo::KanMX* | This study |

**Table S1. Yeast strains used in this study**
